# A cationic motif in Engrailed-2 homeoprotein controls its internalization via selective cell-surface glycosaminoglycans interactions

**DOI:** 10.1101/2021.07.29.454375

**Authors:** Sébastien Cardon, Gérard Bolbach, Yadira P. Hervis, Chrystel Lopin-Bon, Jean-Claude Jacquinet, Françoise Illien, Astrid Walrant, Delphine Ravault, Bingwei He, Laura Molina, Fabienne Burlina, Olivier Lequin, Alain Joliot, Ludovic Carlier, Sandrine Sagan

## Abstract

Engrailed-2 (En2) is a transcription factor that possesses as most homeoproteins the unique and intriguing property to transfer from cell to cell through unconventional pathways. The internalization mechanism of this cationic protein is far from being fully understood and is proposed to require an initial interaction with cell-surface glycosaminoglycans (GAGs). To decipher the role of GAGs in the recognition of En2 at the cell surface, we have quantified the internalization of the homeodomain region in cell lines that differ in their content in cell-surface GAGs. The binding specificity to GAGs and the influence of this interaction on the structure and dynamics of En2 was also investigated at the amino acid level. Our results show that a high-affinity GAG-binding hexadecapeptide (RKPKKKNPNKEDKRPR) located upstream of the homeodomain controls internalization efficiency of En2 through selective interactions with highly-sulfated GAGs of heparan sulfate type. Our data underline the functional importance of the intrinsically disordered basic region that precedes the prominent internalization domain in En2, and demonstrate the critical role of GAGs as an entry gate for En2, finely tuning its capacity to internalize into cells.

## Introduction

Homeoproteins (HPs) are a large family of transcription factors found in all eukaryotes, which are active during development and in adulthood. Beside regulating gene expression, HPs also act as paracrine factors thanks to their unique ability to travel from cell to cell through unconventional transfer (1). Their paracrine action implies a direct access of the traveling proteins to the cytosol and nucleus of recipient cells. HP atypical paracrine activity relies on common structural features shared by all HPs (2), and includes specific secretion and internalization motifs (3). These motifs reside in the 60-residue DNA-binding homeodomain (HD) that defines the HP family (Fig. 1*A*, *B*). HDs are organized as three stable helices while the *N*- and *C*-terminal ends are variable and mostly unfolded (4). The third helix is responsible for the internalization of the protein while the secretion property requires a motif spanning both second and third helices. Notable is that the internalization motif identified 30-years ago within the drosophila *Antennapedia* HD corresponds to the cationic hexadecapeptide Penetratin (RQIKIWFQNRRMKWKK), which has led to the emergence of the field of cell-penetrating peptides (CPPs) (5–7). Importantly, two modes of internalization co-exist for HPs and CPPs, endocytosis and direct translocation, distinguished by their sensitivity to low temperature (1, 5, 8, 9), and by their ability to direct the protein into the cytosol (10).

**Figure 1.**
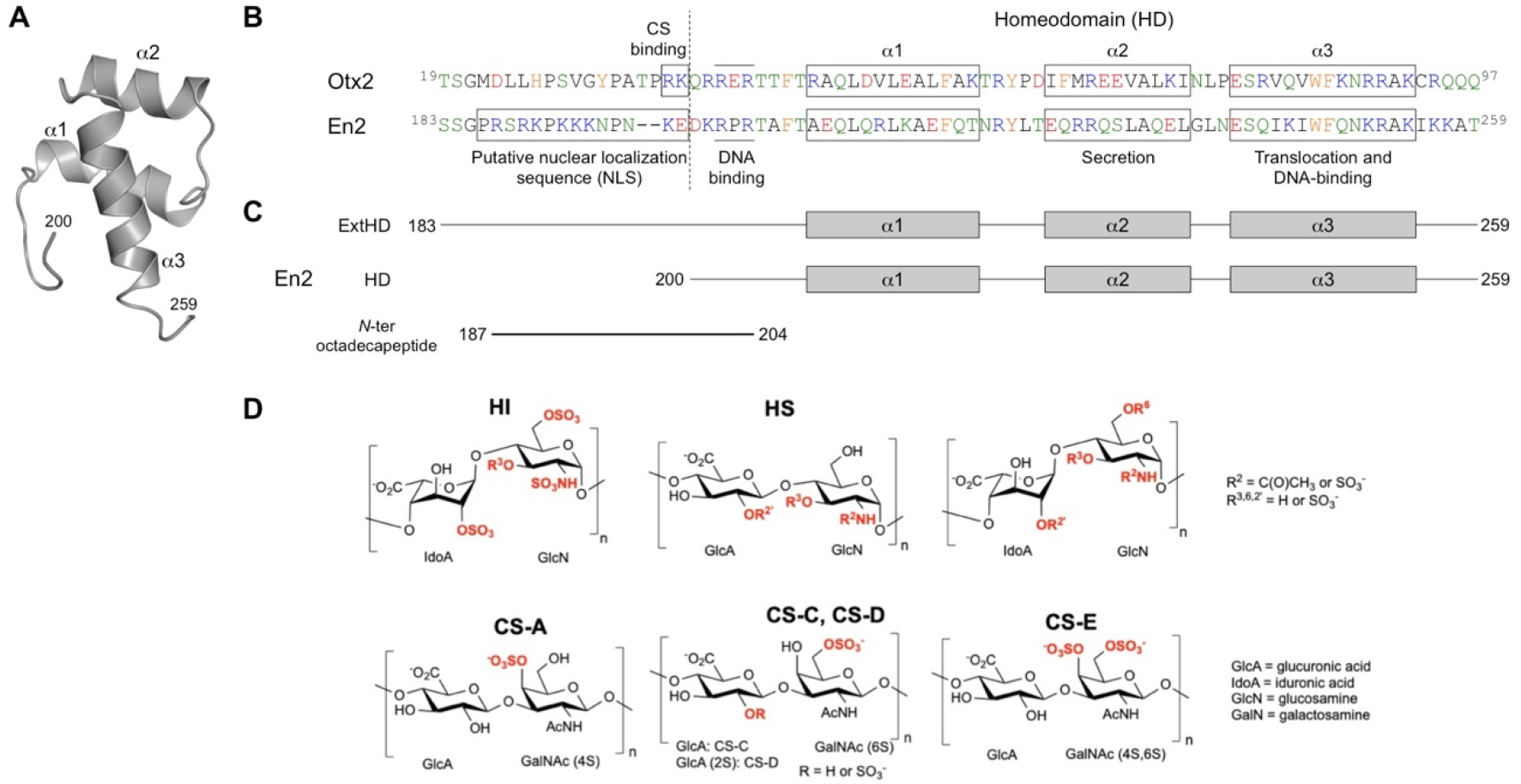
Structure and recognition sequences of En2 region 183-259. **(*A*)** Solution structure of the homeodomain of chicken En2 (region 200-259, PDB code 3ZOB). **(*B*)** Sequence alignment of En2 region 183-259 (chicken gene numbering) with Otx2 region 19-97 (numbering is identical for chicken and mammals). Note that these regions are strongly conserved (99% sequence identity between chicken and human En2, and 100% sequence identity between chicken and human Otx2). Positively charged, negatively charged, polar, apolar, and aromatic amino acids are colored in blue, red, green, black, and orange, respectively. The putative nuclear localization sequence (NLS) of En2 and the RK doublet of Otx2 that was shown to be crucial for the binding to highly sulfated CS (16) are highlighted by rectangles. **(*C*)** Topology of En2 proteins and peptide used in this study. **(*D*)** Structure of disaccharide units of heparan sulfates (HS), chondroitin sulfates (CS), and heparin.

CPP internalization depends on the presence of cell-surface glycosaminoglycans (GAGs) both *in vitro* (8, 11–14) and *in vivo* (15). Regarding HPs, the internalization of extracellular Orthodenticle homeobox 2 protein (Otx2) is restricted *in vivo* to parvalbumin neurons of the developing mouse visual cortex. Secreted Otx2 accumulates in these neurons through interactions with surrounding perineural nets (PNNs) that are enriched in disulfated chondroitin sulfate (CS) GAGs (16, 17). A RK doublet within Otx2 (Fig. 1*B*), upstream of the highly conserved homeodomain, is responsible for the specific interaction with disulfated chondroitin having sulfate group at position 6 (CS-E, CS-D), leading to the restricted internalization of the protein in parvalbumin neurons.

Another homeoprotein, Engrailed-2 (En2), behaves differently. En2 controls the patterning of vertebrate embryos, in particular through the regulation of boundary formation in the developing brain (18, 19). Extracellular En2 does not accumulate in parvalbumin neurons (16) and the region preceding the homeodomain strongly differs from that of Otx2 (Fig. 1*B*). In contrast to Otx2, this region is highly enriched in basic aminoacids and has sequence similarities with nuclear localization signals (NLS).

In this context, we have questioned herein the influence of heparan sulfate (HS) and CS in the internalization, structure, and dynamics of En2. To decipher the role of En2 interactions with GAGs, we have compared the internalization efficacy of two En2 fragments in two cell lines differing in their cell-surface content in CS and HS. Two protein constructs were produced: the homeodomain alone (HD), containing the minimal cell-penetration sequence Penetratin, or extended at its *N*-terminal end with the cluster of basic aminoacids (ExtHD) (Fig. 1*C*). We have characterized at the molecular level the thermodynamics of interaction of both En2 fragments with selected HS and CS molecules, and studied at the amino-acid level, the impact of these interactions on the secondary and tertiary structure of the proteins. We have identified a high-affinity GAG-binding motif located upstream of the homeodomain, which controls the internalization of En2 through selective interactions with highly sulfated HS at the cell surface.

## Results

### The internalization efficacy of ExtHD depends on the presence of HS at the cell surface

First, the uptake efficacies of both En2 fragments were analyzed. A MALDI-TOF-based internalization assay was previously shown to accurately quantify the cellular uptake of biotinylated cell-penetrating peptides (20, 21). We have adapted this protocol to proteins. The method relies on the calibration of the internalized unlabeled protein with the addition of a known amount of its isotopically ^15^N-labeled counterpart just prior cell lysis and before affinity capture through a biotin bait introduced on a cysteinyl residue added at the *N*-terminus (Fig. S1). The quantification is achieved through the ratio of the respective integrated peak area of the non-labeled and ^15^N labeled protein, here used as internal standard. The rate of labeling was calculated from the ion signals (m/z) to be >90%. Despite this high rate of labeling, the ^14^N and ^15^N protein ions have overlapped m/z signals in MALDI-TOF MS spectra. To overcome this overlapping, we developed a software allowing the deconvolution of the signals and an accurate fit of the ratio between the ^14^N non-labeled and ^15^N labeled species (details are included in *SI Appendix*). By comparing the experimental mass spectra to those obtained from the mixing of the individual spectra respectively for the labeled and the non-labeled protein, we were able to determine the absolute quantity of protein internalized inside cells (Fig. S2).

The two En2 fragments were tested for their internalization efficiency in two ovarian cell lines that differ in their content of cell-surface glycosaminoglycans. CHO-K1 cells contain HS and mono-sulfated CS (in a ratio about 50:50) (22, 23). The modified cell line CHO-pgsA-745 (GAG^deficient^) has a genetic deficiency in xylosyl transferase and expresses 10- to 30-fold less HS and CS compared to the parent K1 cell line (22, 23). In K1 cells, HD internalized at very low levels compared to ExtHD (Fig. 2*A*). After one-hour incubation with each protein added at 7 μM, intracellular HD concentration reached 0.13 μM (assuming a total volume of 1 μL for 10^6^ cells). In contrast, ExtHD was internalized 10-fold better. In GAG-deficient cells, internalization of ExtHD was decreased by 6-fold, while HD internalization was only decreased by 2-fold. These results demonstrate that the presence of the *N*-terminal basic extension in ExtHD significantly increases the efficacy of internalization through a process that depends on the presence of cell-surface GAGs. To further analyze the respective contributions of HS and CS on ExtHD internalization, we used different mixtures of enzymes to selectively deplete cell-surface glycosaminoglycans in K1 cells.

**Figure 2.**
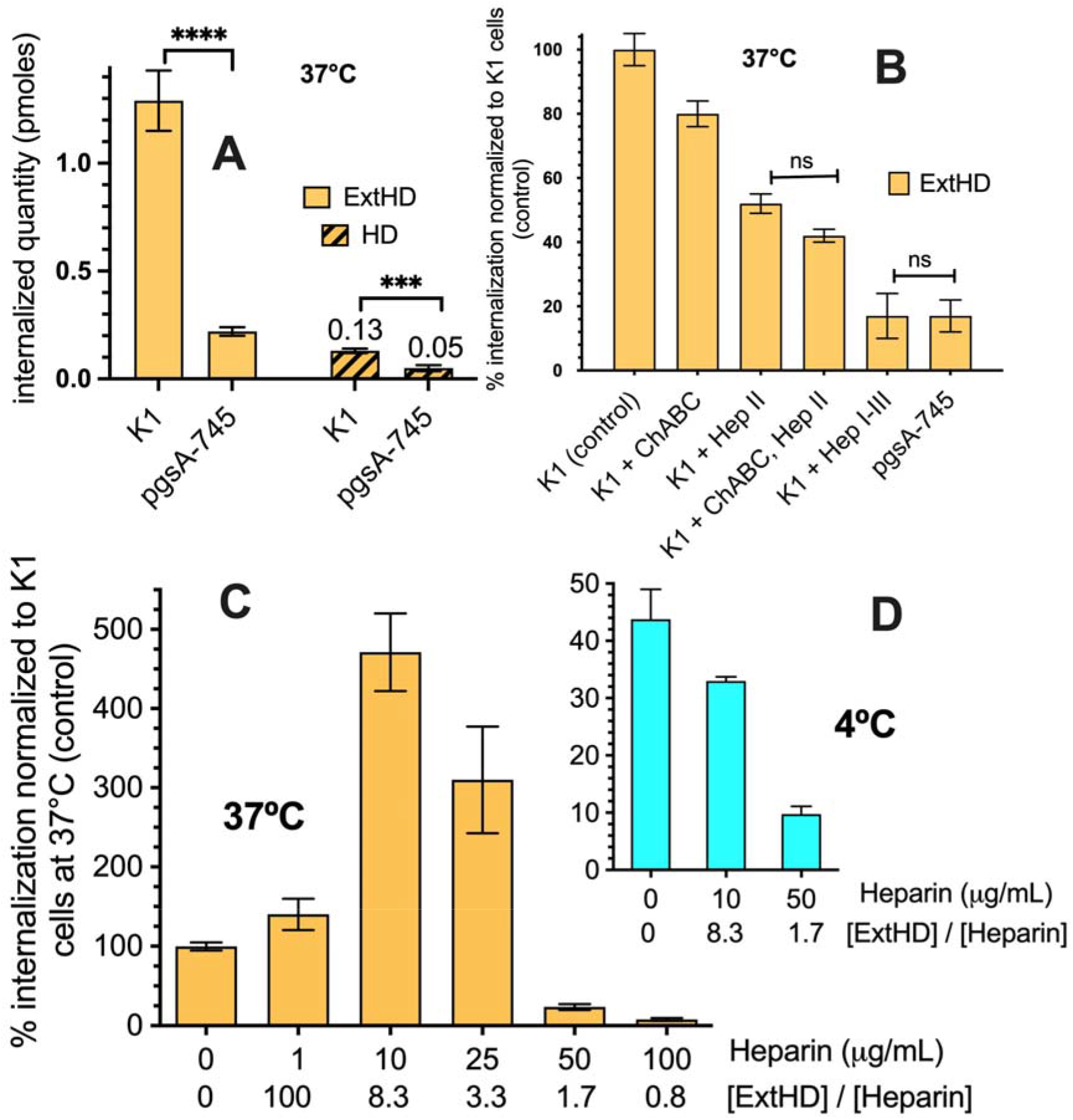
**(*A*)** Quantity of internalized ExtHD and HD incubated at 7 μM with cells for 1 hr at 37°C. CHO-K1 cells express similar levels of HS and mono-sulfated chondroitins; GAG^deficient^ CHO cells (pgsA-745) express 10-30 fold lower GAGs levels than CHO-K1 (22). **(*B*)** Quantity of internalized ExtHD incubated at 7 μM with cells for 1hr at 37°C. CHO-K1 cells (that express HS and mono-sulfated chondroitins) (24) were treated with enzymes degrading HS (heparinases I, II, III) or CS (chondroitinase ABC). Data were normalized to the quantity of internalized ExtHD in control CHO-K1 cells. One-way ANOVA and Bonferroni’s Multiple Comparison Test were used to compare all columns, two by two: only not significant differences (ns) are indicated, all other relevant comparisons being significant (Table S1). **(*C*, *D*)** Quantity of internalized ExtHD in CHO-K1 cells in the absence and presence of increasing amounts of extracellular heparin. One million cells were incubated with 7 μM protein in DMEM-F12 medium for 1 hr either at 37°C **(*C*)** or 4°C **(*D*)**. Data were normalized to the intracellular quantity of ExtHD in control CHO-K1 cells at 37°C in the absence of heparin. The ExtHD/heparin molar ratio is indicated for each heparin concentration.

Internalization of ExtHD was slightly sensitive to chondroitinase ABC (ChABC) treatment (Fig. 2*B*). By contrast, heparinase II (HepII) treatment decreased protein internalization by 50%. Combining HepII and ChABC treatments further decreased ExtHD internalization only by 10%. Finally, when cells were treated with mixed heparinases I-III, ExtHD internalization efficacy dramatically dropped to 15% compared to untreated cells, down to levels similar to the ones measured in GAG-deficient cells. Taken together, these results show that the presence of the 17-amino acid extension rich in basic residues that precedes the homeodomain strongly stimulates ExtHD internalization provided that HS are present at the cell-surface, while the GAG-dependence of HD internalization appears less pronounced. We then asked whether specific interactions between the *N*-terminal extension and HS motifs could account for the greater internalization of ExtHD compared to the shorter HD protein.

### ExtHD recognizes only highly sulfated HS motifs

At the cell surface, HS is a heterogeneous linear polymer that contains highly sulfated regions (so-called S-domains) interspersed with less sulfated and nonsulfated regions. To identify if specific structures within HS were responsible for protein binding, we performed a microarray analysis to probe the interaction of ExtHD with 52 structurally defined HS oligosaccharides of different lengths (from 5-mer to 18-mer) and sulfation patterns (Fig. S4). As shown in Figures S5 and S6, the oligosaccharides that bind more efficiently ExtHD are the 12-mer compounds **18** and **19**, which have the greatest number of sulfates and the highest charge densities (*i.e*. number of sulfates per disaccharide unit).

Notable is the presence of two fully sulfated disaccharide units (*N*-, 2-*O*, 3-*O* and 6-*O* sulfation) in compound **18** and one in compound **19**. A significant fluorescent signal was also observed for the 12-mer compound **22** that contains 4 trisulfated disaccharide units lacking 3-*O* sulfation. By contrast, the 12-mer compounds containing only one or two sulfate groups per disaccharide unit (compounds **23** to **25**) did not yield significant signals, regardless of sulfate distribution. Investigating oligosaccharide length, only low-intensity signals were observed for the sulfated compounds of the 6-, 7- and 9-mer series, including the highly sulfated compound **16** (6-mer, 8 sulfates), indicating that ExtHD better recognizes longer HS fragments (≥ 12 saccharide residues). Overall, this screening profile shows that ExtHD binds selectively to long and highly charged HS, with a strong preference for trisulfated and tetrasulfated disaccharide units.

### Soluble HS discriminates between the two internalization modes of ExtHD

To further characterize the role of HS in ExtHD internalization, the effect of exogenous heparin addition was analyzed on ExtHD internalization. The pharmaceutical heparin used in this study is a highly sulfated HS fragment of ~12 kDa, mostly composed of trisulfated GlcNS,6S–IdoA2S but that also contains less common tetrasulfated GlcNS,3S,6S-IdoA2S disaccharide units (Fig. 1*D*). As shown in Figure 2*C*, heparin exhibited bimodal concentration-dependent effects at 37°C, either promoting or inhibiting ExtHD internalization. At concentrations lower than 25 μg/mL (protein/heparin molar ratio > 3), heparin increased intracellular ExtHD up to 5-fold while concentrations above 50 μg/mL (protein/heparin molar ratio < 2) resulted in a substantial decrease of ExtHD internalization, almost to complete inhibition at 100 μg/mL. Interestingly, analysis by dynamic light scattering (DLS) of the mixed heparin/ExtHD solutions (Fig. S7) revealed the formation of aggregated species (up to 40%) of μm size at low heparin concentration (10 μg/mL), but neither with the protein alone nor when heparin and ExtHD were in stoichiometric quantities. Formation of GAG/protein aggregates is known to promote endocytosis of Penetratin-like CPPs (13, 14, 25) but since Penetratin and HDs also enter cells by direct translocation (8, 26, 27), we analyzed the effect of decreasing temperature on ExtHD internalization efficacy in CHO-K1 cells.

At low temperature (4°C), endocytosis pathways are strongly inhibited, while direct crossing of the lipid bilayer, which is not strictly energy-dependent, still occurs. In the absence of exogenous heparin, ExtHD internalized in CHO-K1 at 4°C with 44 ± 6% efficacy compared to 37°C control condition (Fig. 2*D*), confirming that the protein efficiently internalizes through direct translocation in cells. Contrasting with 37°C conditions, at 4°C the presence of 10 μg/mL heparin in the culture medium resulted in the inhibition of ExtHD internalization in K1 cells (Fig. 2*D*). We concluded that low concentration of soluble heparin can fulfill, even outperform, the requirement of membrane-associated forms of HS for ExtHD endocytosis likely through formation of large complexes between heparin and ExtHD. Conversely, translocation would specifically require interaction with membrane-associated HS, which is antagonized by soluble heparin. Corroborating these results, ExtHD translocation (at 4°C) was impaired in GAG-deficient CHO psgA-745 cells that have reduced levels of membrane-bound HS (Fig. S3). When increasing soluble heparin concentration to reach a protein/heparin molar ratio below 2, aggregates were no longer formed and only competition effects were observed for the two modes of internalization.

### The high binding affinity of ExtHD for heparin relies on the cationic *N*-terminal extension

To further understand the molecular basis of En2 recognition by highly sulfated HS, we used ITC to determine the thermodynamic parameters of heparin interaction with HD and ExtHD proteins. The two protein fragments differed in their affinities for heparin, ExtHD exhibiting a 20-fold greater affinity than HD (Table 1 and Fig. S8).

**Table 1.** Interaction thermodynamics of the *N*-ter octadecapeptide and proteins with heparin, studied by ITC. The proteins were titrated with the polysaccharide at 25°C in 50 mM NaH_2_PO_4_, pH 7.4; nd, not determined.

| polysacch.<br>(z-) | protein<br>(z+, z-) | NaCl<br>(mM) | K<sub>d</sub> (nM) | ΔH<br>(kJ/mol) | -TΔS<br>(kJ/mol) | n<br>(prot/pol<br>y-sacch.) | Complex<br>net<br>charge |
| --- | --- | --- | --- | --- | --- | --- | --- |
| Heparin<br>12 kDa<br>(80-) | *N*-ter<br>octadecapeptide<br>(10+, 2-) | 100 | 300 ± 67 | -81 ± 7 | 44 ± 5 | 7 ± 1 | 56+/80- |
|  | HD<br>(14+, 6-) | 0 | 870 ± 120 | -86 ± 10 | 51 ± 10 | 4.3 ± 0.5 | 38+/80- |
|  |  | 100 | 2090 ± 10 | -46 ± 8 | 14 ± 7 | 5.3 ± 0.5 |  |
|  |  | 200 | nd | nd | nd | nd |  |
|  | ExtHD<br>(21+, 7-) | 0 | 50 ± 10 | -91 ± 2 | 50 ± 8 | 3.5 ± 0.5 | 45+/80- |
|  |  | 100 | 120 ± 18 | -41 ± 2 | 1 ± 1 | 2.9 ± 0.6 |  |
|  |  | 200 | 620 ± 130 | -19 ± 2 | -16 ± 2 | 3.4 ± 0.1 |  |

These interactions were enthalpy-driven, showing that electrostatic interactions are involved in the formation of the complex between either protein and heparin. This observation is further supported by the influence of the ionic strength of the buffer in the formation of the complexes. The reduction from 100 mM to 0 mM NaCl increased the binding affinity by two-fold for both proteins. In contrast, at 200 mM NaCl, affinity was no longer measurable for HD and decreased 5-fold for ExtHD compared to the affinity measured in the 100 mM NaCl condition. Interestingly, a synthetic octadecapeptide corresponding to the cationic region that precedes the helical core of En2 homeodomain (aa 187-204; Fig. 1C) had similar affinity for heparin as the entire ExtHD protein, indicating that this motif is the main determinant of HS interaction within ExtHD. The stoichiometry of the complexes, respectively 5 HD and 3 ExtHD per polymer chain of heparin (12 kDa, 20 disaccharides, ~80 negative charges), also supports the major contribution of the *N*-ter octadecapeptide in this interaction. The octadecapeptide represents indeed only 17% in mass of the total protein while its presence accounts for 40% of the protein binding per heparin chain (see below, Table 3). ExtHD can accommodate 6-7 disaccharides (20 disaccharides in heparin / 3 bound proteins) while HD and the *N*-ter octadecapeptide can bind 4 (20/5) and 3 (20/7) disaccharides, respectively. As the stoichiometry (polysaccharide/protein or 1/n determined in ITC experiments) of interaction between ExtHD and heparin corresponds to the sum of the stoichiometry of interaction measured with the *N*-ter octadecapeptide and HD alone respectively, cooperative events in this interaction appear unlikely. Altogether, these results indicate that the *N*-ter octadecapeptide confers a unique property to the ExtHD protein in terms of binding to highly sulfated HS.

### The binding affinity of ExtHD for phospholipids is not enhanced by the cationic *N*-terminal extension

We next investigated whether enhanced affinity with phospholipids also contribute to the higher internalization efficacy of ExtHD. It is indeed well established that protein-lipid interactions play a key role in membrane-translocation properties of homeodomains (10). In particular, we previously showed that the interaction of En2 homeodomain with anionic lipid vesicles induces a major rearrangement of its tertiary structure (4). Herein we used ITC to probe the interaction of HD and ExtHD with 100 nm large unilamellar vesicles (LUVs) that we prepared either from 1-palmitoyl-2-oleoyl-*sn*-glycero-3-phospho-(1’-rac-glycerol) (POPG), 1-palmitoyl-2-oleoyl-*sn*-glycero-3-phosphocholine (POPC), or a combination of both. The presence of one saturated and one unsaturated fatty acyl chains makes these phospholipids good mimics of mammalian cell membrane composition.

With pure zwitterionic lipids (Table 2), no interaction occurred with either of the two proteins used at 20 μM as previously reported for the HD construct (4). Addition of anionic POPG to POPC to obtain LUVs with 50/50 POPG:POPC and 70/30 POPG:POPC composition, had no effect. Significant ExtHD and HD interactions were observed only with pure POPG LUVs, showing that negatively charged partners are mandatory for interaction of the proteins with lipid membranes. The dissociation constant value determined for POPG was the same for the two proteins (≈ 7 μM; Table 2). These interactions with LUVs were slightly endothermic and entropically-driven, suggesting that water and sodium ions are favorably released from the membrane vicinity concomitantly with proteins transfer from a polar to non-polar environment. Thus, the two proteins have similar binding thermodynamics for POPG, and, importantly, the presence of the cationic *N*-terminal extension does not strengthen the interaction of ExtHD with anionic phospholipids.

**Table 2.** Thermodynamics of interaction of ExtHD and HD with POPG LUVs (100 nm), determined by ITC. The proteins were titrated by LUVs at 25°C in 50 mM NaH_2_PO_4_ buffer, pH 7.5.

| Protein | $K_d^{\text{apparent}}$<br>($\mu\text{M}$) | $\Delta H$<br>(kJ/mol) | $-T\Delta S$<br>(kJ/mol) | n (lipids /<br>protein) |
| --- | --- | --- | --- | --- |
| HD | $7.5 \pm 2.5$ | $1.6 \pm 0.1$ | $-30 \pm 5$ | $7.6 \pm 0.8$ |
| ExtHD | $6.8 \pm 2.3$ | $3.1 \pm 1.3$ | $-29 \pm 3$ | $8.7 \pm 2.3$ |

### ExtHD contains two GAG-binding sites with different affinity for heparin oligosaccharides

We next investigated the molecular basis of the interaction between En2 and highly sulfated GAGs at the single residue level using heteronuclear single-quantum correlation (HSQC) NMR experiments. For that purpose, we expressed and purified ^15^N- and ^15^N/^13^C-labeled HD and ExtHD and obtained the site-specific assignment of backbone resonances. To identify GAG-binding regions of ExtHD, we performed chemical shift perturbation (CSP) experiments by collecting ^1^H-^15^N HSQC spectra on uniformly ^15^N-labeled samples in the presence of increasing amounts of unlabeled oligosaccharides. Our initial attempts to titrate ^15^N-labeled ExtHD with 12 kDa heparin resulted in dramatic line broadening of NMR signals (Fig. S10), in agreement with the formation of a high molecular weight complex as determined by ITC (3:1 protein:GAG complex of 40.5 kDa). Since both the glycan microarray analysis and ITC indicated that ExtHD can accommodate oligosaccharide fragments containing ~6-7 trisulfated disaccharides, we selected a heparin hexadecasaccharide (dp16) fragment for NMR titrations. However, solubility issues were systematically encountered during titration experiments with heparin dp16, and high-quality spectra were only obtained with dp8 and smaller heparin oligosaccharides, although we expected weaker affinities for heparin fragments containing less than 12 monosaccharides. As shown in Figure 3*A*, the addition of stoichiometric amounts of heparin dp8 induced large chemical shift changes in the ^1^H-^15^N HSQC of ExtHD, consistent with a fast exchange regime on the chemical shift time scale and a dissociation constant in the micromolar-millimolar range.

**Figure 3.**
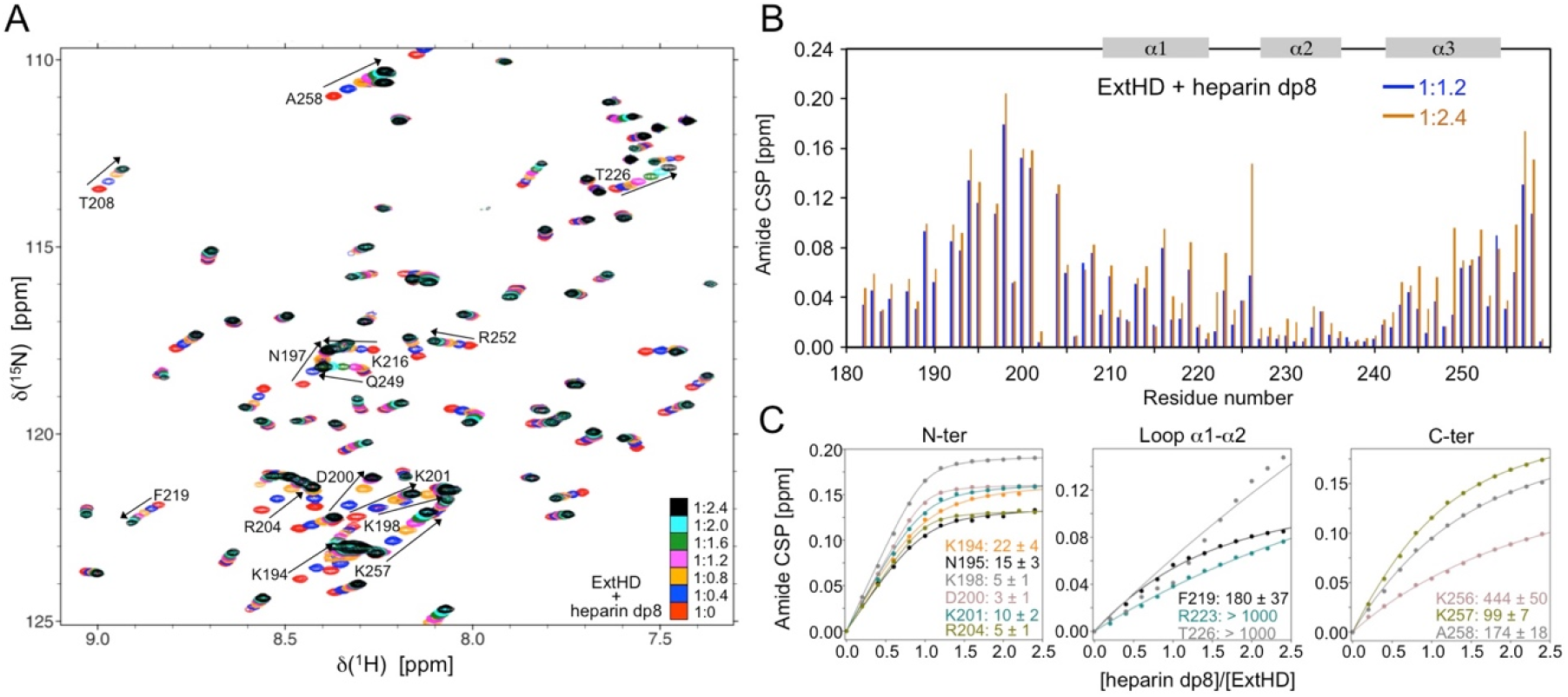
Interaction of ExtHD with heparin dp8 probed by NMR in 50 mM NaH_2_PO_4_ (pH 7.4) and 100 mM NaCl. (***A***) Overlay of ^1^H-^15^N HSQC spectra (500 MHz) obtained for ^15^N-ExtHD in the absence and presence of increasing amounts of heparin dp8. Residues exhibiting the highest perturbations are indicated in the spectra. (***B***) Amide CSP of ExtHD induced by the addition of 1.2 and 2.4 molar equivalents of heparin dp8. (***C***) Saturation curves corresponding to the binding to heparin dp8 are shown for residues displaying the highest perturbations with experimental data represented by points and non-linear curve fitting by lines. Kd^app^ values are given in micromolar.

The highest perturbations were mostly localized in the cationic region ^189^RKPKKKNPNKEDKRPR^204^, which includes the basic residues preceding helix α1 known to interact with DNA (Fig. 3*B*). Large CSP values were also observed at the *C*-terminal tail of the homeodomain that encompasses three positively charged residues (^254^KIKKAT^259^). By contrast, residues of the helical domain displayed rather modest chemical shift changes in the presence of 1.2 molar equivalent of heparin dp8. Additional large perturbations are nevertheless observed beyond a 1:2 protein:GAG ratio in helix α1 (K216 and F219), the loop α1-α2 (R223 and T226), and helix α3 (Q249, N250, K251, and R252). Overall, the high content of basic residues in the most perturbed regions of ExtHD confirms the importance of electrostatic interactions for heparin binding.

In the frame of a binding model involving a two-state equilibrium in the fast exchange regime, the evolution of the ^1^H-^15^N chemical shifts with the protein/oligosaccharide molar ratio can be fitted to determine apparent thermodynamic affinities (K_d_^app^) at the residue level. Variable K_d_^app^ values in the low micromolar range (~5-20 μM) were obtained in the *N*-terminal region, the highest binding affinities being observed in the basic epitope 189-204 (Fig. 3*C* and Table S3). The *C*-terminal residues 256-258 exhibit K_d_^app^ values in the range of 100-400 μM and bound heparin dp8 with a significantly lower affinity. In the case of residues R223 and T226, the binding curves were characteristic of very weak interactions with K_d_^app^ values in the millimolar range. As a result, the loop α1-α2 did not contribute significantly to the binding to heparin despite the large CSP values measured in this region above a protein:GAG ratio of 1:2. These additional CSPs are likely due to a conformational rearrangement of the loop α1-α2 that could be induced by its close spatial proximity with the *C*-terminal heparin binding site or supplemental transient contacts with the heparin fragment.

Taken together, our data reveal that the helical core of En2 homeodomain is flanked by two heparin-binding motifs with residues 192-204 forming the major binding site, and the *C*-terminal residues 254-259 forming an additional, low-affinity binding region. To confirm that these two motifs do not bind heparin in a synergistic manner, we titrated heparin dp8 at increasing concentrations into a solution of ^15^N-labeled HD. The absence of residues 183-199 in the HD construct does not modify significantly the perturbation profile of residues 200-259, which CSP values were remarkably similar to the ones measured with the longer ExtHD construct (Fig. S11). This observation is consistent with the existence of independent interacting regions that bind heparin with differential affinities.

To identify basic residues within the *N*-terminal region that predominantly contribute to heparin binding, we produced three ExtHD mutants in which three basic pairs in the region ^189^<u>RK</u>PK<u>KK</u>NPNKED<u>KR</u>PR^204^ were individually replaced by serine pairs. ITC data revealed that all three mutants lose affinity for heparin compared to the parent protein, from 5-fold for the R189S/K190S mutant up to 8.5-fold for the K193S/K194S one (Table S2 and Fig. S8). For all mutants, the binding enthalpy of the complexes was slightly decreased compared to the wild-type protein, in agreement with the partial loss of electrostatic interactions between these ExtHD mutants and heparin. This decreased enthalpy was partly compensated by a more favorable entropy, likely resulting from increased degrees of freedom for the mutant complexes. Finally, the two mutants K193S/K194S and K201S/R202S exhibited the lowest binding affinities, in the μM range, confirming that basic residues within the region 192-204 are mostly responsible for the binding to heparin.

### The *N*-terminal GAG-binding motif of ExtHD remains disordered upon binding to heparin oligosaccharides

We next investigated the influence of heparin binding on the conformational properties of En2 region 183-259. For that purpose, we examined the site-specific secondary structure of ExtHD in the absence and presence of heparin dp8 using ^1^Hα chemical shift deviations (CSDs). CSDs are defined as the differences between measured chemical shifts and corresponding random coil values for each residue (28). Previous studies have shown that residues 183-206 upstream of the homeodomain are highly disordered in an En2 fragment encompassing residues 146-259 (29, 30). Accordingly, small near-zero values characteristic of disorder were observed in the region 183-206 of ExtHD in the absence of heparin dp8 (Fig. 4*B*). By contrast, the three α-helices of the homeodomain were well identified by three continuous stretches of negative CSDs. The addition of 2.4 molar equivalents of heparin dp8 did not induce significant changes in the ^1^Hα chemical shifts of ExtHD, and the CSD diagram of the bound form was virtually identical to that of the free form. Together with the fact that the presence of heparin fragments only induced selective perturbations of the ^1^H-^15^N chemical shifts, our data indicate that the binding to heparin dp8 does not modify the secondary and tertiary structure of ExtHD. To verify that the conformational properties of ExtHD do not depend on oligosaccharide chain length, we recorded circular dichroism (CD) spectra in the absence and presence of 1 molar equivalent of 12 kDa heparin.

**Figure 4.**
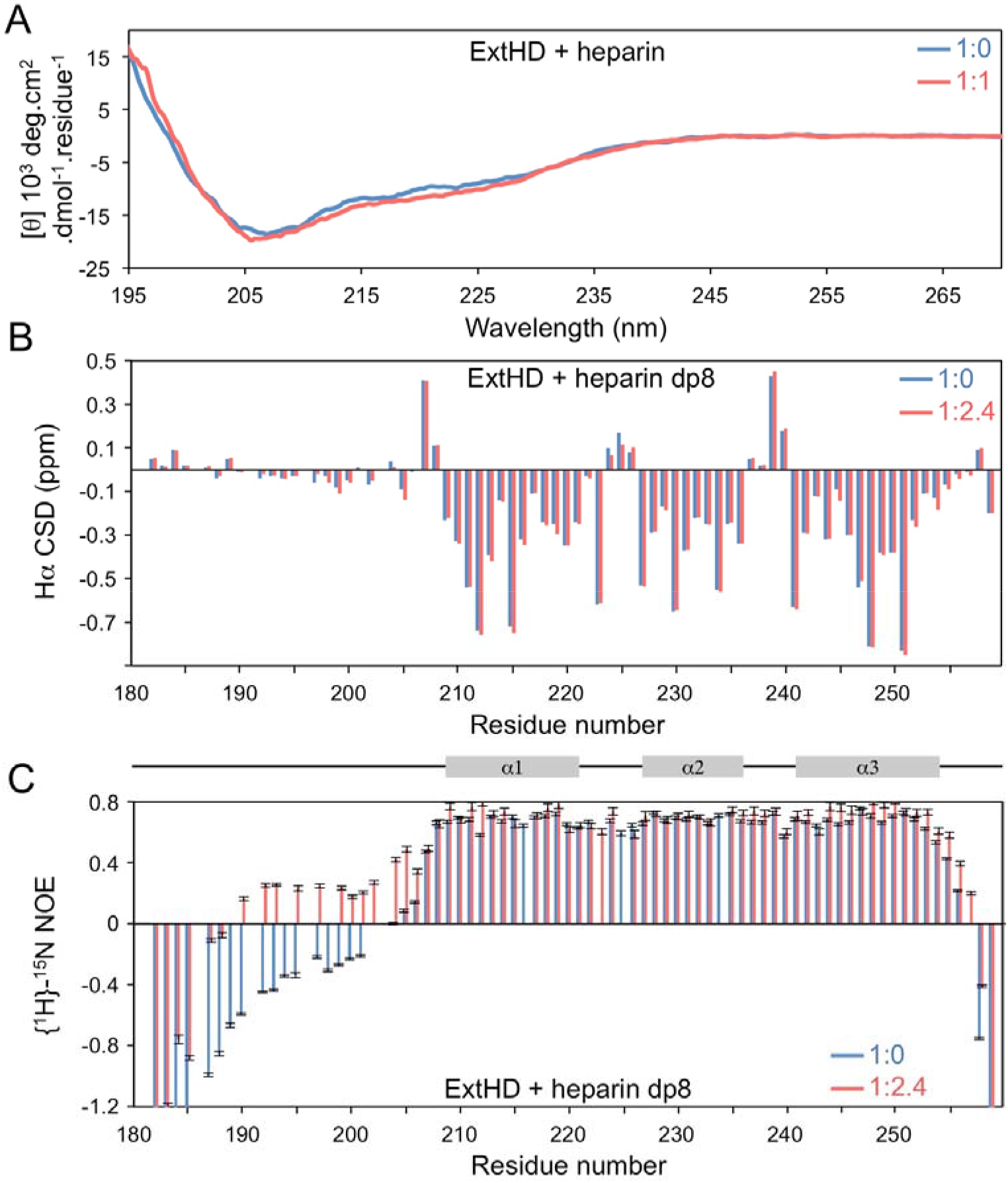
The GAG-binding region 189-204 of Engrailed-2 remains disordered upon binding to heparin or heparin fragments. (***A***) CD spectra of ExtHD in the absence and presence of 1 molar equivalent 12 kDa heparin. Chemical shift deviations (CSD) and {^1^H}-^15^N NOEs of ExtHD recorded at 500 MHz in the absence and presence of 2.4 molar equivalents heparin dp8 are shown in panels (***B***) and (***C***), respectively.

The CD spectrum of ExtHD free in solution displayed two negative ellipticity minima near 208 and 220 nm that are consistent with the α-helical fold adopted by the homeodomain (Fig. 4*A*). The presence of heparin did not modify significantly the CD spectrum, confirming that ExtHD, and in particular the *N*-terminal residues 192-204 do not undergo major structural rearrangement upon GAG binding. We measured heteronuclear {^1^H}-^15^N nuclear Overhauser effects (NOEs) to probe backbone motions of ExtHD that occur on the picosecond-to-nanosecond timescale (Fig. 4*C*). In the free form, the disordered nature of the *N*- and *C*-terminal residues flanking the helical core of the homeodomain was confirmed by negative NOEs that are indicative of high flexibility. Upon binding to heparin dp8, NOE values were found to increase in the *N*-terminal region 190-205 and at the *C*-terminal extremity, evidencing that the interaction with GAGs significantly restricts internal motions in these regions. However, the amplitude of NOEs in the bound form remained weak in the *N*-terminal region (< 0.25) when compared to the rigid helical core that exhibits NOEs > 0.7. As a result, the backbone of the *N*-terminal GAG-binding site retained substantial mobility when bound to heparin, which supports a binding mode mostly driven by electrostatic contacts.

### ExtHD selectively interacts with heparin compared to disulfated chondroitin

As evidenced above, ExtHD preferentially interacts with highly sulfated regions of heparan sulfates. Since other GAG types can harbor high sulfate content, such as sulfated chondroitin (CS) enriched in the PNNs of developing cortex, we investigated whether ExtHD also recognizes structurally defined CS oligosaccharides. As shown in Fig. 1*D*, a number of differences distinguish the molecular structure of CS from that of heparin, including the nature of the amino sugar in the disaccharide repeats (galactosamine in CS versus glucosamine in heparin), glycosidic linkages, the lack of *N*-sulfation in CS, and importantly, the number and distribution of sulfate groups. The NMR titrations of ExtHD with CS-A and CS-C dp6 fragments (1 sulfate group per disaccharide unit) did not result in significant modifications of protein chemical shifts. By contrast, the CSP pattern induced by CS-E dp6 (2 sulfate groups per disaccharide unit) showed strong similarities with that obtained with heparin dp6 (Fig. S12). Large CSP were observed in both the *N*- and *C*-terminal regions flanking the helical domain and the loop α1-α2, indicating that *N*-sulfation of the amino sugar is not mandatory for the binding to ExtHD. Similar to what was observed with heparin fragments, the greatest apparent binding affinities were obtained in the *N*-terminal region 192-204 (Fig. S12 and Table S3). These data suggest that heparin and CS-E oligosaccharides share similar ExtHD-binding sites. However, in comparison with heparin dp6, the CSP magnitude and apparent binding affinities are weaker in both the *N*-terminal region and the *C*-terminal extremity.

To confirm the lower affinity of ExtHD for CS-E, we used ITC and titrated HD and ExtHD proteins with a commercially available CS-E fragment of 72-kDa (Fig. S9). Because the size of this polysaccharide differs from that of heparin, i.e. 12 kDa (~20 disaccharides) for heparin and 72 kDa (~135 disaccharides) for CS-E, the binding affinity for each oligosaccharide type was determined by dividing the measured free energy by the number of bound protein molecules per polysaccharide determined by ITC (see Supplementary Material for further explanation). Our results indicate that ExtHD displays a 126-fold greater affinity for heparin than for CS-E (Table 3). In addition, ExtHD has 17-fold better affinity than HD for heparin whereas the two proteins have similar affinity for CS-E. These data confirm that the presence of the cationic *N*-terminal extension confers to the homeodomain a higher selectivity of recognition for trisulfated HS (heparin over CS-E: selectivity of 126-fold for ExtHD versus 8-fold for HD construct).

**Table 3.** Interaction thermodynamics of the *N*-ter octadecapeptide and proteins with heparin and CS-E, studied by ITC. The proteins were titrated with the polysaccharides at 25°C in 50 mM NaH_2_PO_4_, 100 mM NaCl, pH 7.4. The affinity of peptide or protein for the polysaccharide is determined by dividing the free Gibbs energy by *n* (stoichiometry of the complexes) and calculated as Kd = e ^ΔG/RT^.

| Protein | Polysacch. | $\Delta G_n$<br>(kJ/mol) | *n*<br>protein/<br>polysacch. | $\Delta G_n/n$<br>(kJ/mol) | Kd<br>mM |
| --- | --- | --- | --- | --- | --- |
| *N*-ter<br>octadeca-<br>peptide | heparin | -37 | 7 | -5.3 | 117 |
|  | CS-E | -43 | 55 | -0.78 | 730 |
| HD | heparin | -32 | 5 | -6.4 | 75 |
|  | CS-E | -44 | 33 | -1.3 | 591 |
| ExtHD | heparin | -40 | 3 | -13.3 | 4.5 |
|  | CS-E | -41 | 29 | -1.4 | 568 |

## Discussion

The internalization pathways of homeoproteins and of CPPs derived from the third helix of homeodomains still need to be fully characterized (7, 31). Similarly to the requirement of disulfated GAGs for Otx2 internalization demonstrated i*n vivo*, internalization of Penetratin (the highly conserved third α-helical hexadecapeptide of homeodomains) depends on the presence of cell-surface GAGs (8, 12–15, 32, 33). Therefore, one could assume that Penetratin-like CPPs, HDs, and HPs share at least some steps of their mechanisms of internalization (34). Using a combination of quantitative methods (MS, NMR, ITC, CD), the aim of this study was to explore the role of specific cell-surface GAGs in the recognition and internalization process of the homeoprotein Engrailed-2 (En2), and to identify protein motifs involved in the interaction.

Our data show that the ExtHD fragment encompassing En2 residues 183-259 enters cells principally through interaction with glycosaminoglycans of heparan sulfate (HS) type, thanks to the presence of a unique GAG-binding motif ^189^RKPKKKNPNKEDKRPR^204^ that selectively binds highly sulfated HS. Truncation of this high affinity GAG-binding motif that precedes the helical core of the homeodomain drastically reduces (by about 10-fold) its internalization in CHO cells. Beside this high affinity GAG-binding site, NMR studies reveal that the *C*-terminal extremity of the homeodomain, also interacts with heparin, however with about two orders of magnitude less affinity. The two main GAG-binding regions flanking the helical domain interact with heparin independently from each other. Indeed, the stoichiometry of the interaction between ExtHD and heparin determined by ITC, strictly corresponds to the sum of the ones of HD and the *N*-ter octadecapeptide, taken separately. This result confirms the independence and absence of cooperativity of the two binding sites when the protein interacts with heparin.

The high-affinity HS-binding sequence lies in a highly flexible region of the protein (29, 30), which remains disordered upon binding to GAGs. It contains the BBXBX motif (where B is a basic residue) previously described as a consensus HS-binding sequence (35). It also holds the KKK sequence identified as one of the most significant tripeptides (99^th^ percentile) found among 437 heparin-binding proteins (36). Accordingly, mutation of basic pairs within the ^192^K<u>KK</u>^194^ and ^201^<u>KR</u>PRT^205^ sequences lead to a 10-fold reduction in the binding affinity for heparin. Mutation of the basic pair R189/K190 also decreases the affinity for heparin by 5-fold, showing that the full hexadecapeptide region 189-204 is required to maintain high-affinity binding to heparin. We can thus assume that the full HS binding motif of En2 corresponds to the sequence (B)_2_X(B)_3_(X)_3_BAABBXXB where A is an acidic residue (B and X being respectively a basic and any AA). Interestingly, this motif corresponds to an evolutionary conserved region in engrailed genes sequences.

An important feature of cell-surface HS is the high variability in size, composition, and sulfate distribution of the sulfated regions (S-domains) in which disaccharide units can be sulfated at 4 distinct positions. Using a glycan microarray analysis, we showed that En2 region 183-259 binds preferentially to long (> dp12) and highly sulfated HS oligosaccharides. The correlation between binding affinity and HS sulfate density, together with the strong dependence of the interaction on salt concentration supports a charge-based binding mode. Accordingly, the high-affinity HS-binding motif of En2 can also accommodate the highly sulfated polysaccharide CS-E (2 sulfates per disaccharide unit), albeit with a much lower affinity in comparison with the binding to heparin (3 sulfates per disaccharide unit). ExtHD shows indeed a 126-fold greater affinity for heparin than for CS-E, confirming that En2 interaction with GAGs depends mainly on sulfate density exposed to the protein.

A specific feature of the internalization of HP and HP-derived domains (HD, Penetratin) is the co-existence of two mechanisms, endocytosis and translocation. For all, internalization indeed occurs at 4°C while endocytosis is prevented, and mutations that decrease Penetratin uptake efficiency, such as the replacement of the critical Trp residue by Phe, also prevent HD and HP internalization (5, 37). Translocation implies a transient perturbation/disorganization of the lipid bilayer (3, 34, 38, 39) and it has been recently shown in cell culture that direct interactions between En2 and the anionic phospholipid PIP2 promote the translocation of the protein towards the cytosol (10). Moreover, Carlier et *al*. demonstrated that interaction of En2 homeodomain with anionic phospholipids is required to induce a conformational change in its 3D structure, leading to the insertion of the third helix within the acyl chains of the lipid bilayer (4). However, the plasma membrane is a highly asymmetric structure. In particular, negatively charged lipids are under-represented in the outer membrane leaflet facing the extracellular space, raising the issue of the initiation of the translocation event leading to internalization.

Our data herein support a new role of HS as a critical actor of En2 internalization not only through endocytosis (40), but also through direct translocation across the plasma membrane. En2 translocation (measured at 4°C) is reduced in GAG-deficient cells compared to wild-type cells. Based on previous reports (4) and the data in the present work, we propose that sulfated GAGs anchored by proteins at the cell surface would be good candidates as translocation initiators (Fig. 5). In this model, the highly sulfated S-domains of HS attract En2 at the cell-surface through electrostatic interactions with the HS-binding motif that precedes the homeodomain. The dynamic nature of these interactions allows a fast diffusion of En2 within the extracellular matrix leading to its stepwise accumulation at the close vicinity of the plasma membrane. Indeed, the selective inhibitory action of low concentrations of soluble heparin on translocation but not on endocytosis, supports the need for a proximal distance between GAGs and the lipid bilayer to evoke translocation. This initial interaction with HS would bring the homeodomain in close contact to the acyl part of the lipid bilayer (41), leading to the modification of its conformation and the insertion of the third helix within the lipid bilayer. The interaction of En2 with PIP2 in the inner leaflet, shown to regulate the bidirectional exchanges of the protein between the cytosol and the lipid bilayer (10), would eventually result in the transfer of the protein within the cytosol. Following this model, we propose that the bidirectional transfer of En2 is regulated by a dynamic equilibrium between protein-lipid or protein-GAG interactions depending on the side of the plasma membrane.

**Figure 5.**
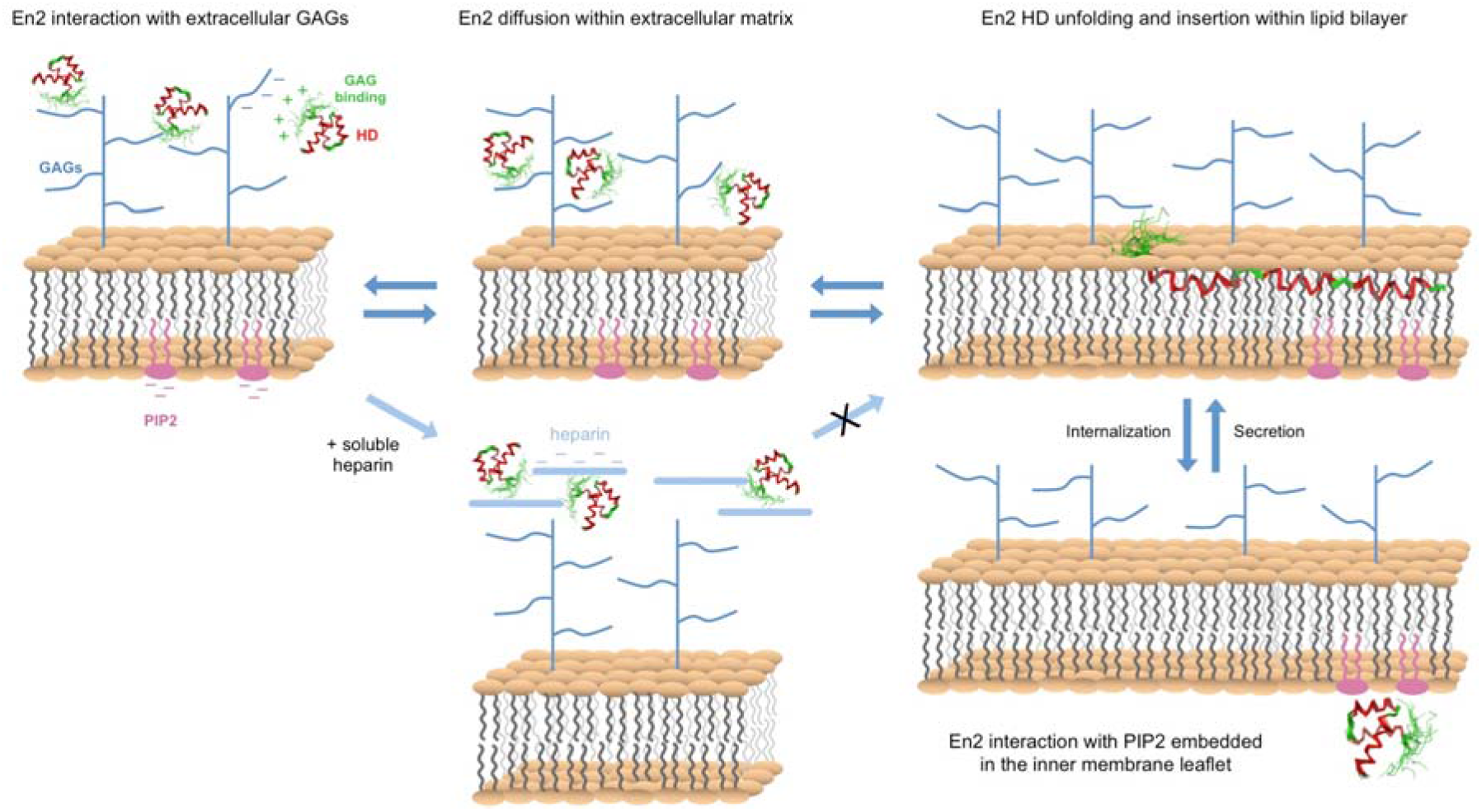
Model of GAG-dependent membrane translocation of HPs (see text for details).

The interaction of GAGs with extracellular HPs has important consequences at the physiological level, as the paracrine action of HPs relies on their ability to be internalized. For instance, the plasticity of the visual cortex relies on the ability of secreted Otx2 to internalize into parvalbumin neurons through interactions with the surrounding extracellular matrix enriched in disulfated GAGs (16, 17). The axonal guidance of retinal neurons also depends on the paracrine action of HPs, Vax1 at the optic chiasma and later, En2 in the tectum (42, 43). Vax1 internalization requires the presence of HS on retinal neurons, and for En2, its delivery into the cytosol is required to exert its paracrine action that involves the translational activation of Ephrin-mediated axon guidance (42, 44). Since a graded distribution of En2 in the extracellular matrix of the tectum is necessary for the proper positioning of retinal axons in the developing brain (45), GAGs might also act to restrict HP diffusion in the extracellular space (46, 47), protecting them from enzymatic degradation (Fig. S13).

A critical component of HP paracrine action is the restriction of their internalization to specific cell types. Due to their widely heterogenous structures, GAGs are attractive candidates to modulate the cellular tropism of HP internalization. Compared to En2, the binding specificity for GAGs is markedly different in the case of Otx2 homeoprotein, which contains only 5 positively charged residues, including a RK doublet, in the *N*-terminal extension preceding the homeodomain helical core (Fig. 1). *In vivo*, an Otx2 fragment encompassing the homeodomain and the RK doublet is selectively internalized into cortical neurons surrounded by perineuronal nets similarly to the full protein, while the corresponding AA double-mutant is not. Mutation of the RK doublet by alanines in an Otx2 pentadecapeptide (^36^<u>RK</u>QRRERTTFTRAQL^50^) fully abolishes peptide binding to CS-D and CS-E oligosaccharides (16). These results show that the RK doublet upstream of the homeodomain is responsible for the specific interaction of Otx2 with cell-surface CS and its subsequent internalization in PNN-positive cells. Interestingly, the GAG-interacting domains within En2 and Otx2 strongly differ, the RK doublet that recognized disulfated CS in Otx2 being replaced by a KE doublet in En2. KE pairs are frequently found in HS-binding proteins (36). In addition, our combined ITC and NMR data show that En2 region 183-259 (ExtHD) interacts more efficiently with heparin than disulfated CS-E, while Otx2 displays 10-fold higher affinity for CS-E and CS-D as compared to heparin (16). Accordingly, En2 is significantly less internalized in PNN-positive cells than Otx2 (16). Overall, these findings support the idea that the cationic motifs preceding the homeodomain region control the spatial distribution of extracellular homeoproteins through a fine tuning of their binding specificity for GAGs. This underlines the importance of these GAG-binding motifs that define a new functional region involved in HP trafficking, acting in concert with internalization/secretion motifs.

## Conclusions

This study highlights the crucial role of GAGs in the control of the internalization of En2 homeoprotein. We provide evidence at the molecular and cellular levels of a direct correlation between the selective binding of En2 protein to highly sulfated HS and its capacity to internalize into cells expressing HS, including through a translocation process across the plasma membrane. It is likely that such GAG-dependent internalization reported for Otx2, Vax1 and now En2 also applies to other homeoproteins. Here we provide evidence of differential interactions of HPs with GAGs, that would govern the cellular tropism of their paracrine action towards distinct cell types, depending on their GAG content. More precisely, expression of specific membrane-surface GAGs and fine tuning of their sulfation content may control the entry of homeoproteins in cells. Multiple modifications may occur in the HS structure, in particular sulfation or epimerization, which could impact many cellular processes such as differentiation, proliferation, and migration (48). In the brain, this could result in an alteration of neurogenesis, neuritogenesis and angiogenesis, and impact plasticity and homeostasis. Based on our study, modulation of homeoprotein paracrine action might be an attractive effector regulated by fine GAG structure modification.

## Materials and Methods

Detailed description of the materials and methods including protein expression and purification, biotin labeling, cell culture, protein internalization assay, protein-GAGs interaction by ITC and NMR, and protein structure and dynamics by CD and NMR is provided in *SI Appendix*.

## Acknowledgments

This research has received funding from the Agence Nationale de la Recherche program (grant N° ANR-BLAN2016-CROSS).

