## Supplemental Information for "A cationic motif in Engrailed-2 homeoprotein controls its internalization via selective cell-surface glycosaminoglycans interactions"

[c]Univ. Orléans, CNRS, ICOA, 45067 Orléans, France.

[d]INSERM U932, Institut Curie Centre de Recherche, PSL Research University, Paris, France

**Keywords**
Homeoproteins, Engrailed-2, protein-GAG interaction, membrane translocation,

***Corresponding Authors**



### Materials and Methods

#### Expression and purification of recombinant proteins

Regions 200-259 (HD) and 183-259 (ExtHD) of chicken Engrailed-2 (Uniprot accession number Q05917) were expressed as $(His)_6$-tagged Cherry fusion proteins using pSCherry1 vectors and the *E. coli* SE1 strain as expression host (Eurogentec, Seraing, Belgium). ExtHD mutants K189S/K190S, K193S/K194S, and K201S/R202S were expressed as $(His)_6$-tagged GST fusion proteins using pGEX-6P plasmids (Genscript, Piscataway, NJ, USA) and the *E. coli* BL21-CodonPlus-RIL expression strain (Agilent Technologies, Santa Clara, CA, USA). A PreScission protease cleavage site (LEVLFQ/GP) was introduced prior to the En2 sequence in all constructs. As a result of PreScission cleavage, final proteins have four extra residues at the *N*-terminus (GPAS). For internalization assays, an additional Cys residue was introduced at the *N*-terminus by site-directed mutagenesis (leading to GPCAS). Unlabeled fusion proteins were produced from cells cultured in LB rich medium. Uniformly labeled $^{15}N$ and $^{15}N,^{13}C$ proteins were overexpressed in M9 minimum media using the procedure developed by Marley *et* al. for preparing high yields of uniformly labeled proteins (1). Protein expression was induced by adding 1 mM isopropyl-1-thio-β-D-galactopyranoside (IPTG) to the cell cultures followed by a 4-hour incubation period at 37°C under agitation. Cells were pelleted by centrifugation (3000 × g for 20 min), resuspended in a 20 mM $NaH_2PO_4$ buffer (pH 7.4) containing 500 mM NaCl, 30 mM imidazole and 1 mM DTT, and lysed by sonication. Soluble fractions containing En2 proteins were separated from cell debris by centrifugation (30000 × g for 40 min at 4°C), and loaded on a Ni-NTA column (GE Healthcare, Chicago, IL, USA). Proteins were purified using IMAC standard protocols with a 30–250 mM imidazole gradient. Eluted fractions were pooled and dialyzed against PBS buffer, and $(His)_6$-tagged PreScission protease was added to cleave fusion proteins. A second step of purification was further performed using Ni-NTA chromatography to remove the cleaved $(His)_6$-tagged fusion partner and the $(His)_6$-tagged PreScission protease. Purity of protein fractions was controlled by SDS-PAGE analysis, and protein identity was checked by MALDI-TOF analysis. Fractions containing pure target proteins were finally concentrated by ultrafiltration using a 3-kDa cutoff membrane (Millipore). The synthetic *N*-ter octadecapeptide used in ITC experiments (biotin sulfone-$G_4$-RSRKPKKKNPNKEDKRPR-$CONH_2$) was synthesized by the Boc strategy, purified by RP-HPLC (purity > 95%) and checked by MALDI-TOF mass spectrometry (expected mass: 2744.5 Da; measured mass: 2744.9 Da).

### Preparation of $^{14}N$, $^{15}N$ biotin-labeled recombinant proteins

Biotin-labeled proteins were used in internalization assays to fish the internalized species out the cell lysates through specific binding to streptavidin-coated magnetic beads. The two recombinant proteins expressed in *E. coli* contain an additional Cys on the *N*-terminal end. Purified proteins were transferred to a buffer containing 50 mM sodium phosphate (pH 7.0), 150 mM NaCl, 10 mM EDTA, and 400 μM TCEP. The protein solution was degassed for 30 min under vacuum before addition of 5 equivalents of maleimide-PEG$_2$-biotin (MW = 525) (Sigma Aldrich, Saint-Louis, MO, USA). The reaction was carried out under inert atmosphere ($N_2$) to prevent oxidation of the maleimide-PEG$_2$-biotin and decrease of the reaction yield. The reaction solution was protected from the light and stirred for 90 min at room temperature. The reaction was then repeated with addition of fresh maleimide-PEG$_2$-biotin solution. Proteins were transferred into a buffer containing 20 mM $NaH_2PO_4$ (pH 7.2), 20 mM NaCl, and stored at -80°C. The yield of protein biotinylation was evaluated to be > 90%, obtained from MALDI-TOF MS analyses (Fig. S1).

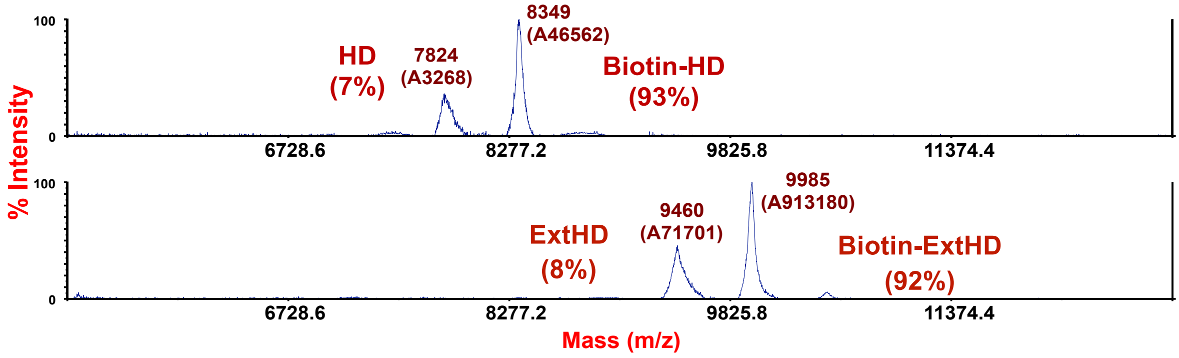

**Fig. S1**. Mass spectra (MH+) of $^{14}N$ biotin-labeled HD and ExtHD proteins. MS spectra were acquired on a Voyager DE-Pro MALDI-TOF apparatus in linear positive mode. The expected masses for biotin-HD and biotin-ExtHD are 8349 and 9985 Da, respectively.

### Cell Culture

Wild type Chinese Hamster Ovary CHO-K1 cells and GAG-deficient CHO-pgsA745 cells (GAG-deficient) (ATCC, LGC Standards S.a.r.l. - France) were cultured in DMEM-F12 medium (Life Technologies, Carlsbad, CA, USA) supplemented with 10% fetal calf serum (FCS), penicillin (100,000 IU/L), streptomycin (100,000 IU/L), and amphotericin B (1 mg/L) in a humidified atmosphere containing 5% $CO_2$ at 37°C.

### Enzymatic removal of glycosaminoglycans from cell-surface

Cells were washed with PBS and incubated with enzymes, alone or in mixture: heparinase I, heparinase II, heparinase III or/and chondroitinase ABC (Sigma Aldrich, Saint-Louis, MO,

USA) at a final concentration at 0.1 U/mL in PBS pH 7.4 (containing calcium and magnesium) for 1 hr at 37°C. Cells were then washed with HBSS, then with DMEM F12 before further incubation with the proteins.

**Quantification of protein internalization by mass spectrometry**

The cellular entry of proteins was quantified by MALDI-TOF mass spectrometry as extensively described for peptides (2), except that we used $^{15}$N-labeled proteins as internal standards and developed a software to calculate the amount of internalized protein. Briefly, adherent and confluent cells ($10^6$ cells/well in 12-wells plates) were incubated with 500 µL of a $^{14}$N-protein in culture medium at 7 µM for 60 min at 37°C. After washing, trypsin treatment of cells permits to digest membrane-bound protein species and to detach cells. Cells are lysed with a hypertonic detergent solution containing a known amount of the $^{15}$N-labeled proteins and boiled. After centrifugation to remove cell debris, the lysate is incubated with streptavidin-coated magnetic beads for 1 hr to extract $^{14}$N- and $^{15}$N- proteins. After several washing steps, proteins are eluted from beads in 3 µL of CHCA matrix at room temperature for 15 min. The samples are then analyzed by MALDI-TOF MS (positive ion linear mode) on a Voyager-DE Pro mass spectrometer (Applied Biosystems, Foster City, CA, USA). Experiments were first done to determine the accurate amount of $^{15}$N-protein to be added to obtain as close as possible a 1:1 ratio between the internalized $^{14}$N and $^{15}$N internal standard proteins. This first step allows one to get accurate, robust and reproducible MS quantification results (3). Each experiment was done in triplicate and independently repeated at least 3 times.

**Software developed to quantify the cellular uptake of proteins by mass spectrometry**

In a previous study (2), MALDI-TOF was found to be well adapted to quantify the cellular uptake of biotinylated cell-penetrating peptides. This method was based on the addition of a known and appropriate amount of the labeled peptide ($^2$H deuterium labeling) to the cell lysis buffer before extraction of the two peptide species ($^2$H-labeled and non-labeled). As in conventional internal standard experiment the quantification is achieved through integrating the peaks area of both labeled and non-labeled peptides. For proteins with higher mass than peptides, as those studied herein, we have developed a software allowing an adjustment of the ratio *r* between non-labeled ($^{14}$N)/labeled ($^{15}$N) proteins, by comparing the experimental mass spectra profile to that obtained from the mixture of separate labeled and non-labeled protein. The validity of such an approach originates from the properties of both proteins: same sequence implying identical ionization yield, similar incorporation in the MALDI matrix, same peak broadening due to attachment of alkaline and matrix adducts, identical detection efficiency due to the small mass difference. In addition, if experimental mass spectra of $^{14}N$, $^{15}N$ and the mixture ($^{14}N$ and $^{15}N$) from internalization are obtained under the same experimental conditions (delayed ion extraction, ion detection and laser fluence), it is possible after mass calibration to calculate a theoretical mass spectrum for a mixture with a ratio *r* and to compare it to the experimental mass spectrum. The software calculates, normalizes and displays the experimental and simulated mass spectra and a fast adjustment of the ratio *r* can be obtained. Baseline correction on the calculated mass spectrum can be also implemented to get a better adjustment simulated vs experimental. Studying the different charge states allows to estimate the accuracy of the ratio.

The non-complete labeling of the protein can be also studied by comparison of $^{14}N$ and $^{15}N$ mass spectra through the calculated convolution of the $^{14}N$ ion peak (+1, +2, +3…) with a Gaussian distribution of $^{15}N$ atoms by adjusting the mean value $<^{15}N>$ and the standard deviation ($\sigma$). The convolution operates over the m/z range in which the $^{14}N$ peak is defined. The display of the convoluted and $^{15}N$ experimental peak allows one to deduce the value of $<^{15}N>$ and $\sigma$. If needed, experimental data can be integrated before processing the convolution.

Experimental MS spectra are extracted as ASCII files (m/z, intensity) from the processing software (Data Explorer) of the MALDI-TOF apparatus (DE-Pro and 4700 Proteomics Analyzer both from Applied Biosystems, Foster City, CA, USA). The software to determine the ratio *r* and also the labeling distribution, if any, has been written in Visual Basic (V 6.0). The simplest case is that of a complete labeling for which non-labeled and labeled peaks are superimposable and thus the m/z shift of the non-labeled peak overlay that of labeled peak using an intensity ratio (Figure S2). To take into account a labeling distribution, each experimental point of the unlabeled peak is shifted in m/z with a relative intensity modulated by the probability of the shift. A new peak is formed after adding all the contribution for a value m/z within a chosen window (1-5 Da depending on the protein mass).

In Fig. S2 an example is shown with the ExtHD protein as unlabeled ($^{14}N$) and labeled ($^{15}N$) versions. No $^{15}N$ distribution was necessary to fit the +1 and +2 protein peaks showing that the labeling was complete. The experimental data of $^{15}N$ were well fitted from the $^{14}N$ data shifted by 128 Da for both +1 (lower mass spectrum) and +2 (upper mass spectrum) charge states.

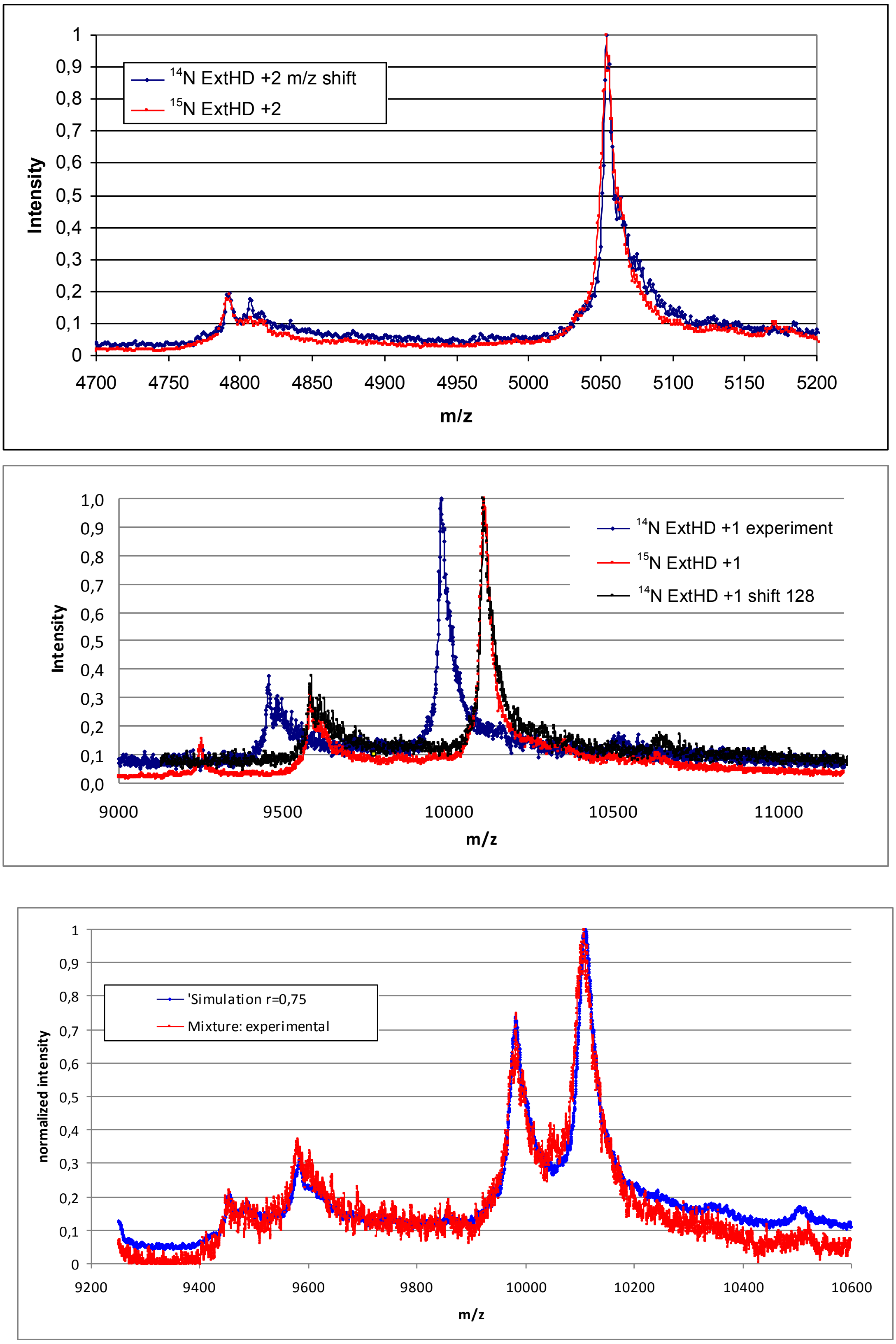

**Fig. S2**. The figure illustrates the fitting of the ratio $r = [^{14}N]/[^{15}N]$ for ExtHD protein using the peak of $^{14}N$ and $^{15}N$ separately recorded in the same conditions and mixed to form a ratio *r* and then compare to the experimental data of the mixture. The best adjustment for the $MH^+$ peak is obtained for a ratio $r = 0.75$. It should be noted that satellites peaks of the protein of interest can also be simulated and indicate a very close ratio *r*.

**Microarray of structurally defined HS compounds**

A custom HS-microarray was designed by the company Glycan Therapeutics (https://www.glycantherapeutics.com/services/microarray-analysis), as previously described (4). 52 structurally defined HS oligosaccharides were synthesized and immobilized on a microarray chip. The biotinylated ExtHD protein was incubated at 100 nM with the microarray slide for 1 hr in a Tris/phosphate buffer pH 7.50 containing 137 mM NaCl and 10% bovin serum albumin (BSA) (4). After washing with the Tris/phosphate buffer, the microarray slide was treated with 10 μg/mL Alexa Fluor 488 labeled Avidin. The wash process was repeated before scanning fluorescence of the array slide at 488 nm.

**Isothermal titration analysis (ITC)**

Experiments were done with a nano-ITC calorimeter (TA Instruments, New Castle, DE, USA). Measurements were done at 25°C in 50 mM $NaH_2PO_4$ pH 7.4 and various NaCl concentrations (0, 100, 200 mM). Titration of the proteins in the ITC cell was done by 25 successive injections of 10 μL of polysaccharide (heparin, heparin-derived, CS-E, obtained from Iduron, Cheshire, UK) with 5 min injection intervals. Proteins, peptide and polysaccharides were used in different concentrations to get accurate fitted curves (between 20-80 μM for proteins and peptide and 40-120 μM for GAG). Control experiments were obtained by injections of buffer into proteins and injections of GAGs into buffer. Data were analyzed with NanoAnalyze software provided by TA Instruments.

LUVs were prepared as previously described (5). Briefly, lipid films were prepared by dissolving phospholipids into chloroform or a mixture of chloroform and methanol (2/1, vol/vol). Formation of the lipid film was achieved by evaporating solvents under nitrogen flux, then drying in a vacuum chamber for at least 1 hr. Films were then hydrated with 10 mM sodium phosphate buffer pH 7.5 and extensively vortexed at a temperature superior to the lipid phase transition temperature to obtain MLVs. To form LUVs, the MLVs were subjected to five freeze/thawing cycles. The homogeneous lipid suspension was then passed 19 times through a mini-extruder (Avanti Polar Lipids, Avanti Alabaster, AL, USA) equipped with two stacked 100 nm polycarbonate membranes at a temperature above the phase transition temperature of the phospholipid. Titrations were performed by injecting aliquots of LUVs (1 mg/mL) into the calorimeter cell containing the protein solution (0.1 mM), with 5 min intervals between injections.

**Circular dichroism (CD)**

CD spectra were measured on a Jasco J-815 spectropolarimeter (JASCO Corporation, Tokyo, Japan) over the wavelength range 190–270 nm, by using a 0.1-cm path-length quartz cell (internal volume 200 μL) from Hellma (Muellheim, Germany). Measurements were carried out at 25°C with a 1 nm/min scan speed and a band width of 1 nm. Proteins and oligosaccharides were dissolved in a buffer containing 20 mM sodium phosphate (pH 7.4) and 100 mM NaF. Four scans were accumulated and averaged for each sample. All spectra were corrected by subtraction of the background obtained for each protein-free mixture. Protein concentration was 10 μM in the absence and presence of 1 molar equivalent heparin.

**NMR data collection and analysis**

NMR experiments were acquired at 298K on a Bruker Avance III 500 MHz spectrometer equipped with a 5-mm TCI cryoprobe. NMR spectra were processed and analyzed with the software programs NMRPIPE (6) and NMRFAM-SPARKY (7). All experiments were performed with NMR samples containing $^{15}N$ or $^{15}N$,$^{13}C$ uniformly labeled HD or ExtHD, previously dissolved in a buffer containing 40 mM sodium phosphate (pH 6.3), 100 mM NaCl, and 7.5% $D_2O$. 0.1 mM Sodium 2,2-dimethyl-2-silapentane-5-sulfonate-d6 (DSS) was added in the NMR samples as an internal $^{1}H$ chemical shift reference. $^{13}C$ and $^{15}N$ chemical shifts were referenced indirectly to DSS, using the absolute frequency ratios. Sequence-specific assignments of backbone $^{15}N^{H}$, $^{1}H^{N}$, $^{13}C\alpha$, $^{1}H\alpha$, $^{13}CO$, and side-chain $^{13}C\beta$, $^{1}H\beta$ resonances were obtained from the analysis of a series of three-dimensional triple resonance experiments (HNCA, HNCO, HN(CA)CO, HNCACB, CBCA(CO)NH, HNHA, and HBHA(CO)NH), recorded on samples containing 600 μM $^{15}N$,$^{13}C$ ExtHD. Chemical shift perturbation experiments (CSP) were performed by collecting $^{1}H$-$^{15}N$ HSQC spectra on $^{15}N$ uniformly labeled HD or ExtHD (100-150 μM) in the absence and presence of increasing amounts of heparin fragments (obtained from Iduron, Cheshire, UK) or structurally defined CS-E dp6 that was chemically synthesized (8), previously dissolved in the NMR buffer. CSP values were calculated for each residue from the differences in $^{1}H^{N}$ ($\Delta\delta_{H^N}$) and $^{15}N$ ($\Delta\delta_{N}$) chemical shifts between the free and the bound states using the relation: $CSP = \sqrt{(\Delta\delta_{H^N})^2 + (0.1\Delta\delta_N)^2}$. Apparent thermodynamic affinities ($K_d^{app}$) at the residue level were calculated for residues in the fast exchange regime from a nonlinear least squares curve fitting of CSPs upon oligosaccharide addition using in-house Python scripts, as previously described (9). Steady state $^{15}N$-{$^{1}H$} NOE values in the absence and presence of heparin dp8 (Iduron, Cheshire, UK) were determined as the ratio of peak heights in $^{15}N$-$^{1}H$ spectra collected with and without proton saturation during the recycle delay. $^{1}H$ saturation was achieved by applying a train of 120° pulses spaced at 5 ms interval for a period of 4.5 s. Uncertainty in the $^{15}N$-{$^{1}H$} NOE values were calculated from the root mean square baseline noise of both spectra estimated with NMRPIPE.

**Quantification of protein internalization by Mass Spectrometry**

**Table S1.** Statistical analysis for significance between means of data presented in Fig. 2 (Quantity of internalized ExtHD or HD incubated at 7 µM with cells for 1 hr at 37°C). Data were analyzed by One-way ANOVA to compare the means of the different cell treatment conditions, followed by a Bonferroni's Multiple Comparison Test to compare pairs of columns (two by two comparison, starting from left to right).

| Parameter | | | |
|---|---|---|---|
| Table Analyzed | Cells + enzymatic treatments | | |
| One-way analysis of variance | | | |
| P value | < 0.0001 | | |
| P value summary | *** | | |
| Are means signif. different? (P < 0.05) | Yes | | |
| Number of groups | 6 | | |
| F | 94 | | |
| R squared | 0,92 | | |
| Bartlett's test for equal variances | | | |
| Bartlett's statistic (corrected) | 70 | | |
| P value | < 0.0001 | | |
| P value summary | *** | | |
| Do the variances differ signif. (P < 0.05) | Yes | | |
| ANOVA Table | SS | df | MS |
| Treatment (between columns) | 99000 | 9 | 11000 |
| Residual (within columns) | 8200 | 70 | 120 |
| Total | 110000 | 79 | |

| Bonferroni's Multiple Comparison Test | Mean Diff. | t | Significant? P < 0.05? | Summary | 95% CI of diff |
|---|---|---|---|---|---|
| K1 (control) vs K1 + ChABC | 20 | 3.9 | Yes | ** | 2.6 to 37 |
| K1 (control) vs K1 + Hep II | 48 | 8.4 | Yes | *** | 29 to 67 |
| K1 (control) vs K1 + ChABC, Hep II | 58 | 11 | Yes | *** | 41 to 75 |
| K1 (control) vs K1 + Hep I-III | 83 | 16 | Yes | *** | 66 to 100 |
| K1 (control) vs pgsA-745 | 83 | 15 | Yes | *** | 64 to 100 |
| K1 + ChABC vs K1 + Hep II | 28 | 4.9 | Yes | *** | 8.6 to 47 |
| K1 + ChABC vs K1 + ChABC, Hep II | 38 | 7.4 | Yes | *** | 21 to 55 |
| K1 + ChABC vs K1 + Hep I-III | 63 | 12 | Yes | *** | 46 to 80 |
| K1 + ChABC vs pgsA-745 | 63 | 11 | Yes | *** | 44 to 82 |
| K1 + Hep II vs K1 + ChABC, Hep II | 10 | 1.8 | No | ns | -9.4 to 29 |
| K1 + Hep II vs K1 + Hep I-III | 35 | 6.1 | Yes | *** | 16 to 54 |
| K1 + Hep II vs pgsA-745 | 35 | 5.6 | Yes | *** | 14 to 56 |
| K1 + ChABC, Hep II vs K1 + Hep I-III | 25 | 4.9 | Yes | *** | 7.6 to 42 |
| K1 + ChABC, Hep II vs pgsA-745 | 25 | 4.4 | Yes | ** | 5.6 to 44 |
| K1 + Hep I-III vs pgsA-745 | 0.00 | 0.00 | No | ns | -19 to 19 |

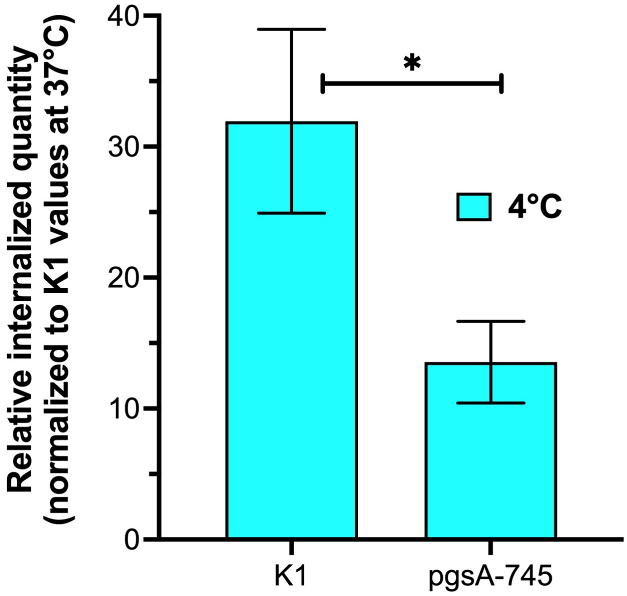

**Fig. S3**. Relative internalization of ExtHD at 4°C in K1 and pgsA-745 cells.

### Screening of ExtHD binding to HS oligosaccharides using a microarray analysis

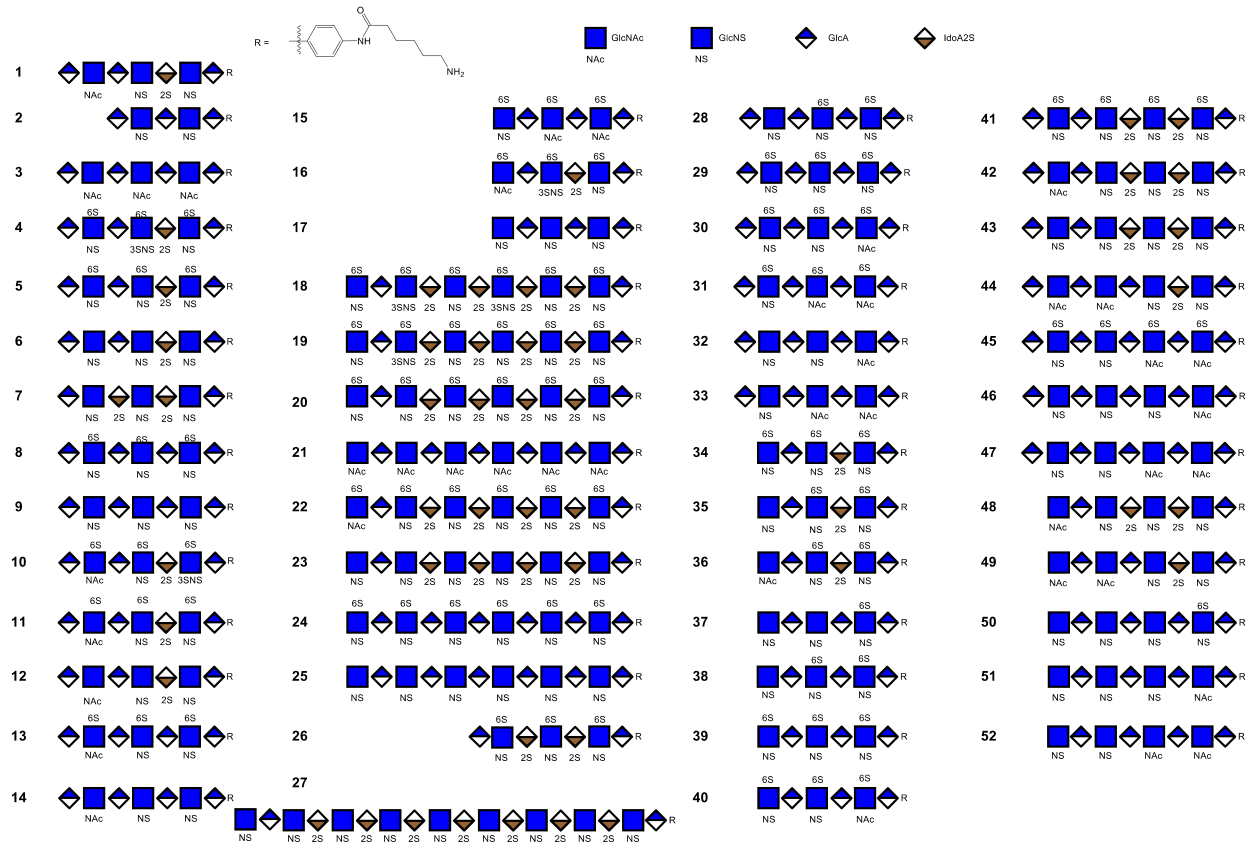

**Fig. S4**. Structurally defined heparan sulfate oligosaccharides used in the microarray analysis to probe the binding to ExtHD (https://www.glycantherapeutics.com/services/microarray-analysis). The 52 compounds exhibit different sulfation patterns usually found in nature, with a size range from 5 (5-mer) to 18 (18-mer) monosaccharide units.

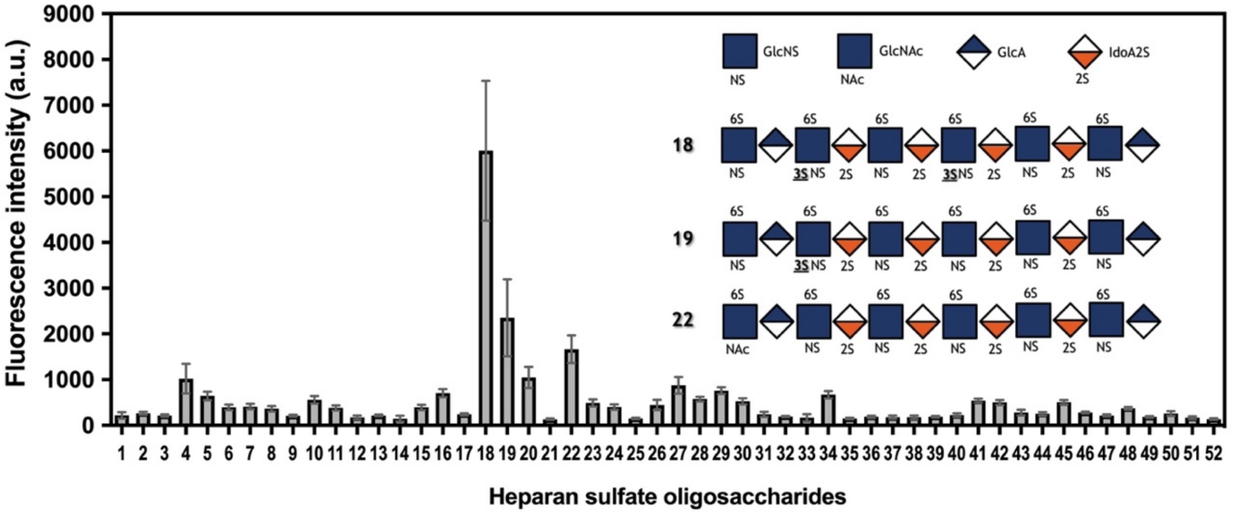

**Fig. S5**. Heparan sulfate microarray analysis. Biotin-labeled ExtHD (100 nM) was incubated on the microarray with different HS structures and its binding revealed by fluorescent streptavidin.

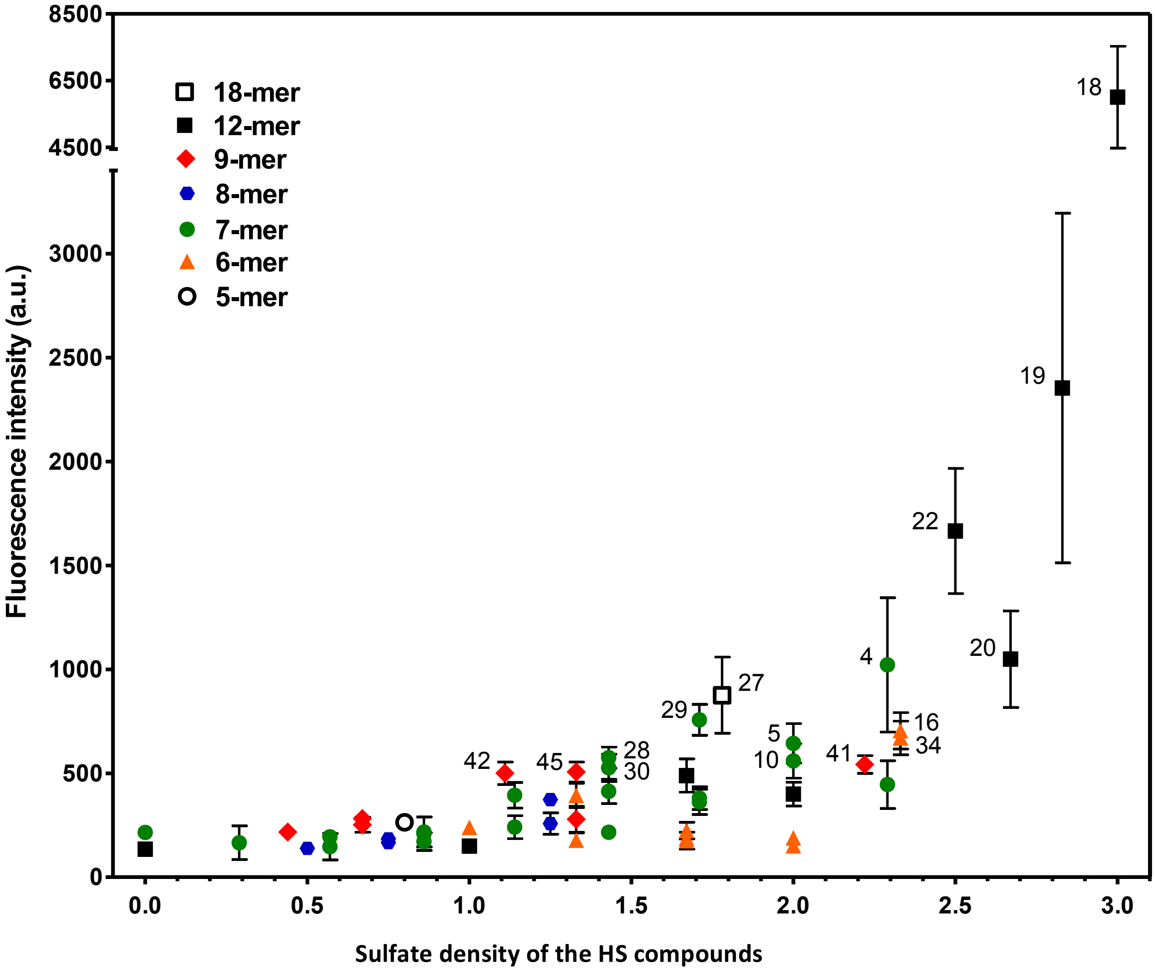

**Fig. S6.** Scatter plot of the measured fluorescence intensity vs. sulfate density (number of sulfates per disaccharide unit) of the 52 HS oligosaccharides used in the microarray analysis to probe ExtHD binding. Compound numbers are indicated for HS oligosaccharides showing fluorescence intensity higher than 500.

### Dynamic light scattering (DLS) analysis of heparin/ExtHD complexes

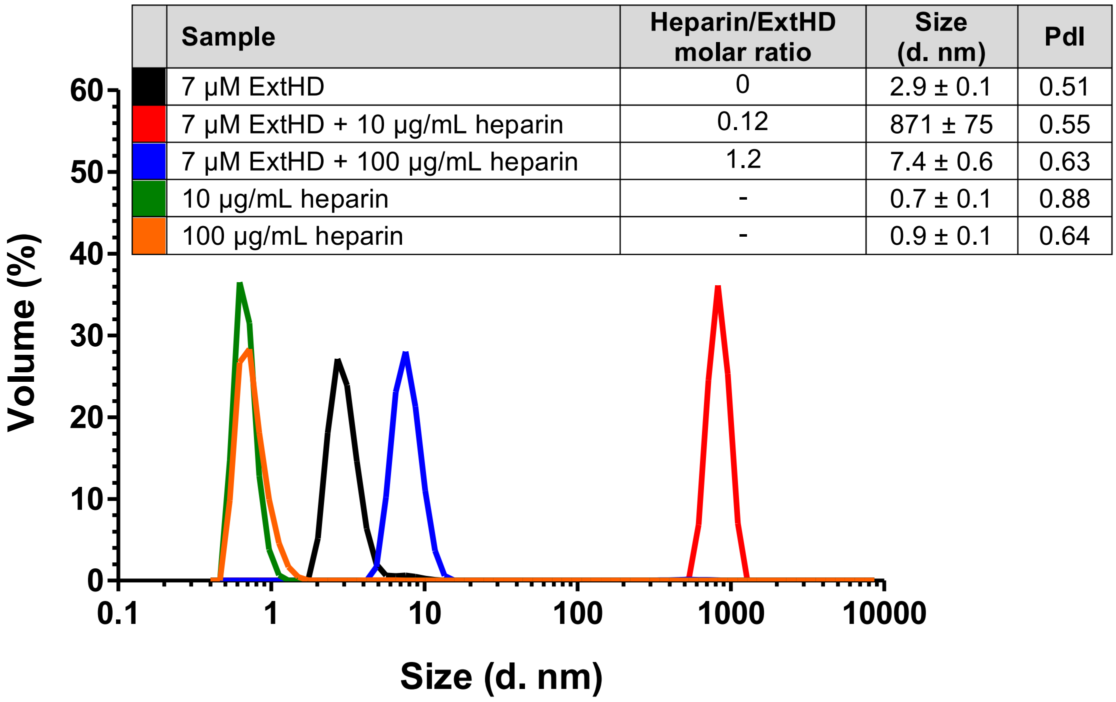

| | Sample | Heparin/ExtHD molar ratio | Size (d. nm) | PdI |
|---|---|---|---|---|
| (black) | 7 µM ExtHD | 0 | 2.9 ± 0.1 | 0.51 |
| (red) | 7 µM ExtHD + 10 µg/mL heparin | 0.12 | 871 ± 75 | 0.55 |
| (blue) | 7 µM ExtHD + 100 µg/mL heparin | 1.2 | 7.4 ± 0.6 | 0.63 |
| (green) | 10 µg/mL heparin | - | 0.7 ± 0.1 | 0.88 |
| (orange) | 100 µg/mL heparin | - | 0.9 ± 0.1 | 0.64 |

**Fig. S7.** Size distribution of ExtHD mixed with heparin. The sample mix was incubated 1 hr at 37°C before recording the mass distribution using a Zetasizer Nano ZS apparatus (Malvern Instruments, Malvern, UK). The table shows the heparin/ExtHD molar ratio for each sample, the size in nanometer, and the polydispersity index (PdI) of particle populations.

### Interaction of En2 proteins with GAGs by Isothermal Titration Calorimetry (ITC)

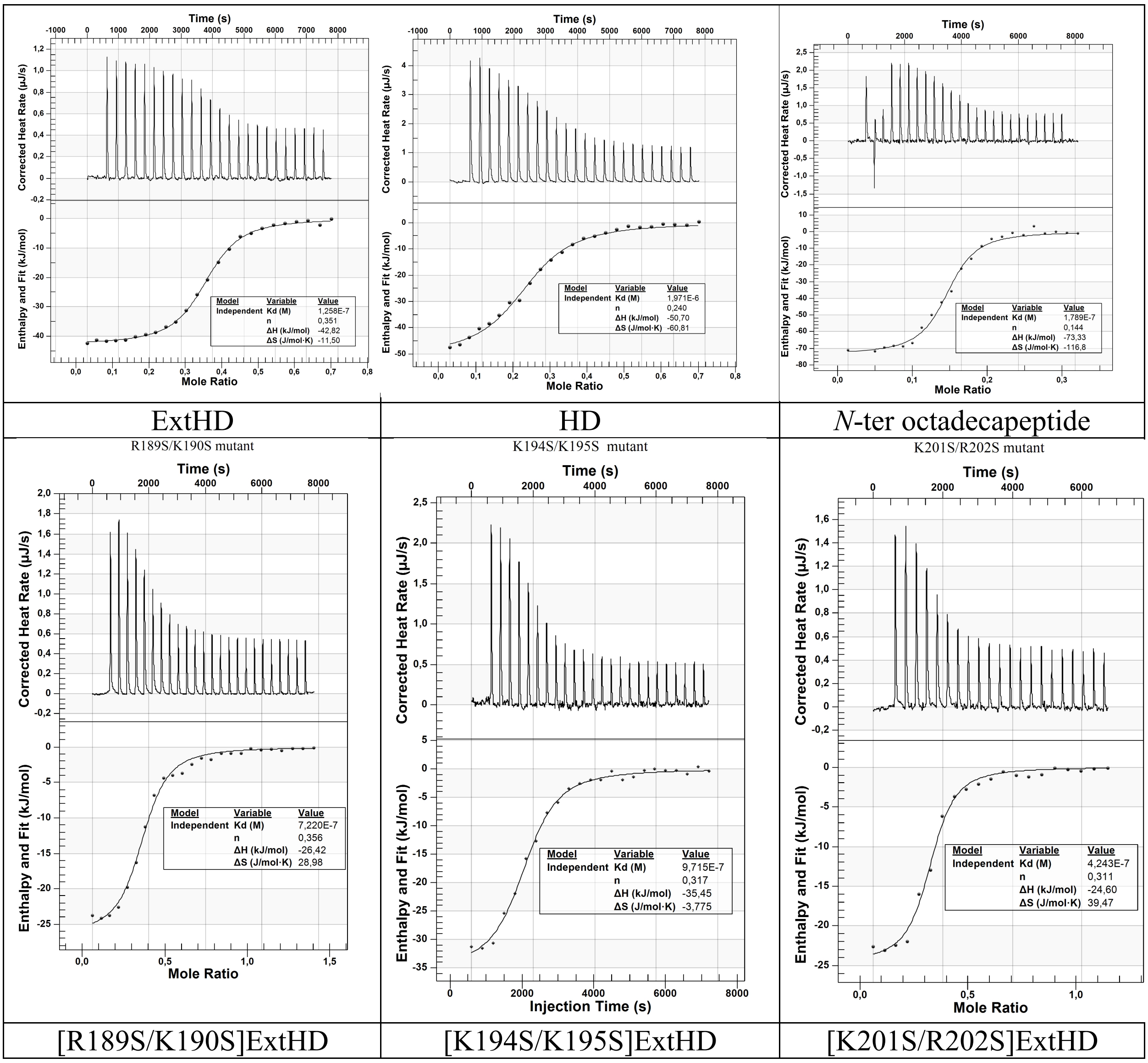

**Fig. S8**. Representative ITC thermograms for the titration of ExtHD, mutants, HD and *N*-ter octadecapeptide with heparin in 50 mM $NaH_2PO_4$, 100 mM NaCl, pH 7.4 at 25°C. Experiments were acquired with a Nano ITC calorimeter (TA Instruments, New Castle, DE, USA) and analyzed with the Nanoanalyze software provided by the supplier.

**Table S2.** Interaction thermodynamics of ExtHD proteins, mutated in the region 189-204 (RKPKKKNPNKEDKRPR), with 12 kDa heparin studied by isothermal titration calorimetry (ITC). The proteins were titrated with the polysaccharide at 25°C in 50 mM $NaH_2PO_4$ (pH 7.4), 100 mM NaCl.

| Protein | Kd (nM) | ΔH (kJ / mol) | -TΔS (kJ / mol) | n (prot/polysacch.) |
|---|---|---|---|---|
| ExtHD | 120 ± 18 | -41 ± 2 | -1.0 ± 1.0 | 2.9 ± 0.6 |
| K193S/K194S | 1010 ± 200 | -34 ± 1 | 0.7 ± 0.1 | 3.0 ± 0.5 |
| K201S/R202S | 925 ± 175 | -28 ± 1 | -6.0 ± 1.0 | 3.3 ± 0.3 |
| R189S/K190S | 628 ± 6 | -26 ± 1 | -9.0 ± 1.0 | 2.7 ± 0.2 |

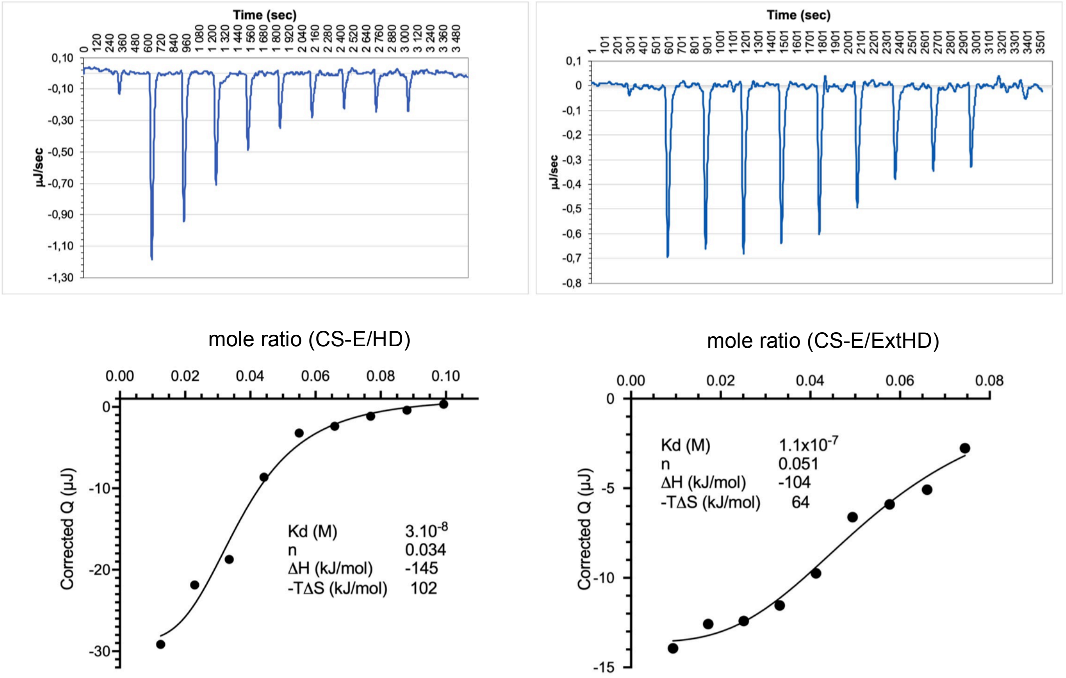

**Fig. S9**. Representative ITC thermograms for the titration of HD and ExtHD with CS-E in 50 mM $NaH_2PO_4$, 100 mM NaCl, pH 7.4 at 25°C. Experiments were acquired with a Nano ITC calorimeter (TA Instruments, New Castle, DE, USA) and analyzed with the Nanoanalyze software provided by the supplier.

**Analysis of the thermodynamic parameters of the interactions of En2 proteins with the CS-E GAG type by ITC (Fig. S9).** Because the size of this oligosaccharide differs from that of heparin, i.e. 12 kDa (~20 disaccharides) for heparin and 72 kDa (~135 disaccharides) for CS-E, we could only compare the energetics of binding for one particular polysaccharide chain, but cannot evaluate accurately the relative affinity of each protein for the two polysaccharides. The polyelectrolyte character of the polysaccharides (their length) is indeed a crucial parameter that influences the binding energetics of the protein. The increase of binding affinity with the length and charge density of these polyelectrolytes is expected owing to increased electrostatic interactions and Coulombic end effects (10) as previously reported for the formation of FGF-2-heparin (11) or Tat-heparin (12) complexes. Hence, to get access to the binding energy of only one protein molecule to one polysaccharide chain, the global free energy of binding was divided by the number of protein molecules bound per polysaccharide (stoichiometry determined in ITC experiments), but with the assumption or approximation that there is no cooperativity between proteins during binding to the polyelectrolyte and that all binding sites are similar.

### Investigation of the GAG-binding properties of En2 proteins by NMR

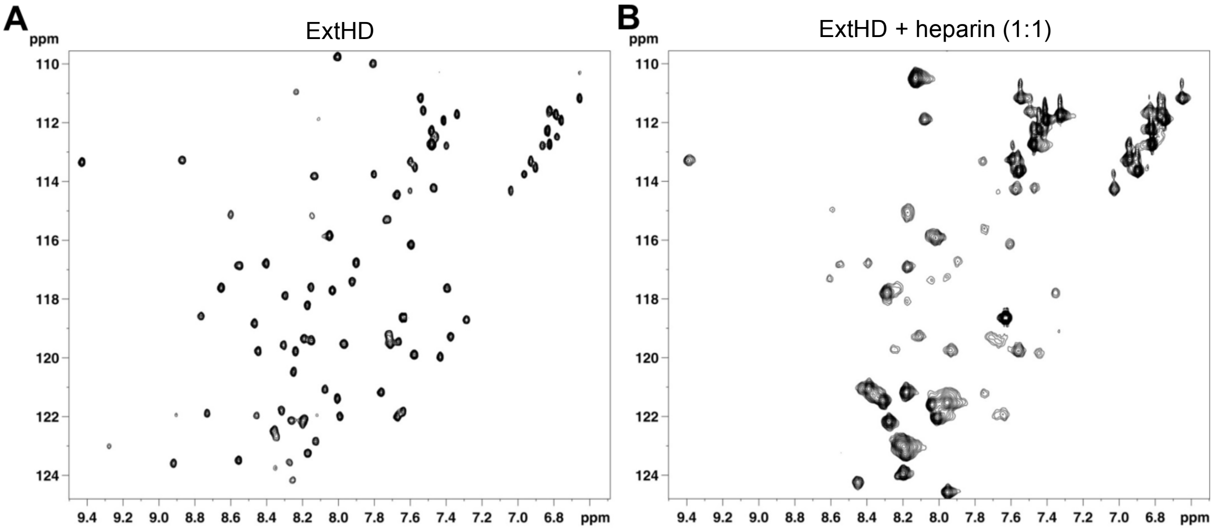

**Fig. S10**. $^{1}H$-$^{15}N$ HSQC of 150 μM $^{15}N$-ExtHD in the absence (*A*) and presence (*B*) of one molar equivalent of unlabeled heparin.

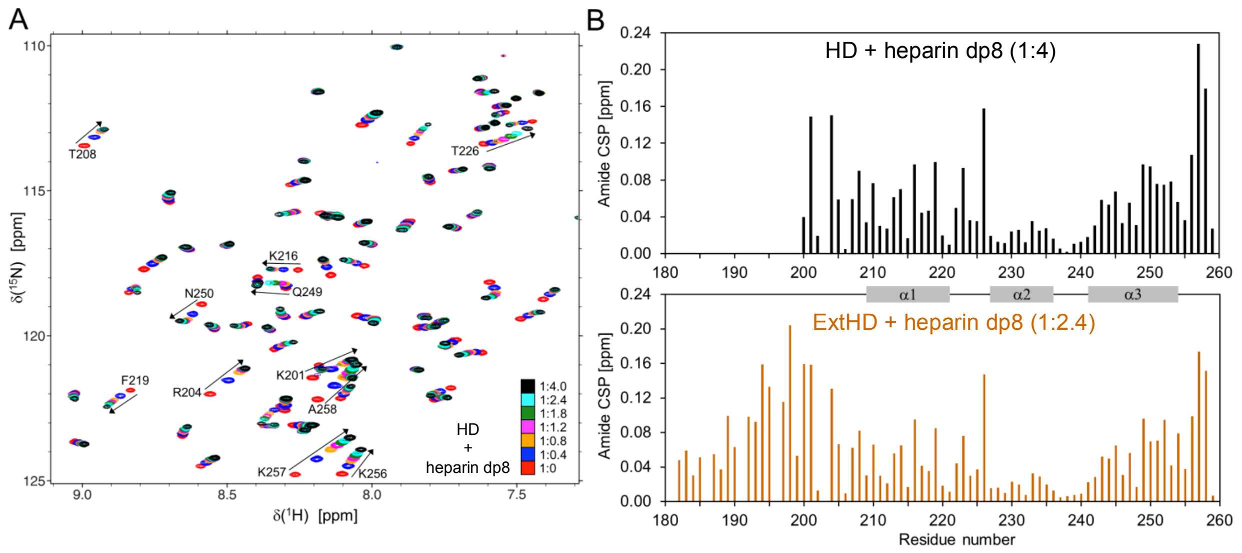

**Fig. S11**. Interaction of HD with heparin dp8 probed by NMR. (*A*) Overlay of $^{1}$H-$^{15}$N HSQC spectra obtained for $^{15}$N-HD in the absence and presence of increasing amounts of heparin dp8. Residues exhibiting the highest perturbations are indicated in the spectra. (*B*) Average amide CSP of HD induced by the addition of 4 molar equivalents of heparin dp8. The CSP profile of ExtHD with heparin dp8 is shown for comparison.

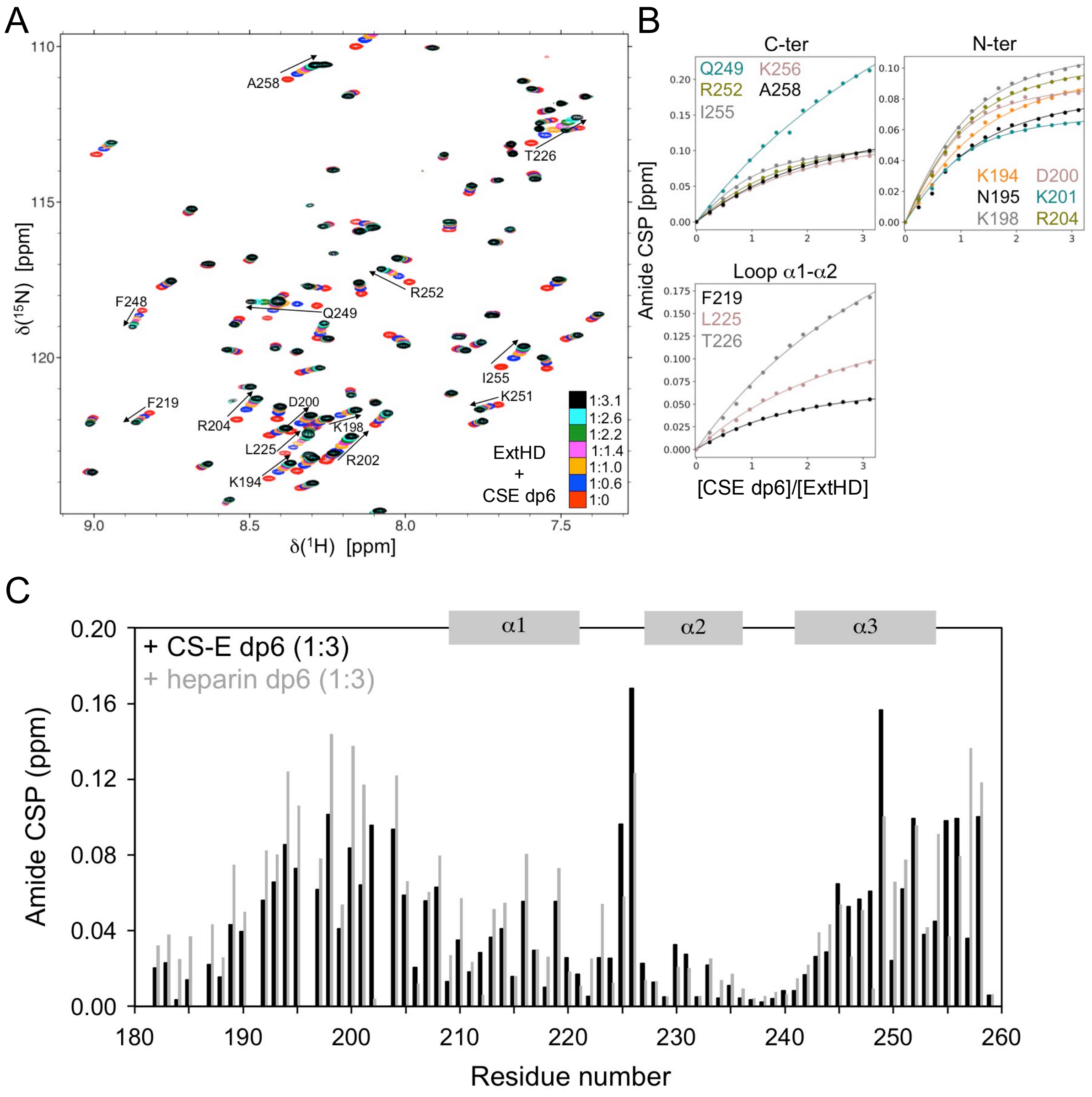

**Fig. S12**. Interaction of ExtHD with structurally defined CS-E dp6 probed by NMR. (*A*) Overlay of $^{1}H$-$^{15}N$ HSQC spectra obtained for $^{15}N$-ExtHD in the absence and presence of increasing amounts of CSE dp6. Residues exhibiting the highest perturbations are indicated in the spectra. (*B*) Saturation curves corresponding to the binding to CSE dp6 are shown for residues displaying the highest perturbations, with experimental data represented by points and non-linear curve fitting by lines. (*C*) Amide chemical shift perturbations (CSP) of ExtHD induced by the presence of saturating amounts of CS-E dp6. For comparison, CSP values obtained with heparin dp6 are shown in grey.

**Table S3**. Apparent $K_d$ ($K_d^{app}$) at the residue level obtained from the non-linear curve fitting of NMR CSPs as a function of oligosaccharide/protein molar ratio. Values are given for representative residues of ExtHD in micromolar.

| Residue | heparin dp8 | heparin dp6 | CSE dp6 |
|---|---|---|---|
| **R189** | 23 ± 5 | 99 ± 11 | – |
| **K192** | 16 ± 3 | 52 ± 8 | 106 ± 17 |
| **K193** | 24 ± 4 | 74 ± 8 | 113 ±15 |
| **K194** | 22 ± 4 | 72 ± 7 | 113 ± 16 |
| **N195** | 15 ± 3 | 46 ± 7 | 99 ± 21 |
| **N197** | 5 ± 1 | – | 29 ± 15 |
| **K198** | 5 ± 1 | 31 ± 4 | 67 ± 8 |
| **D200** | 3 ± 1 | 14 ± 3 | 37 ± 5 |
| **K201** | 10 ± 2 | 35 ± 6 | 49 ± 6 |
| **R204** | 5 ± 1 | 23 ± 3 | 65 ± 8 |

| Residue | heparin dp8 | heparin dp6 | CSE dp6 |
|---|---|---|---|
| **K216** | 3 ± 1 | 18 ± 3 | 61 ± 7 |
| **F219** | 180 ± 37 | 183 ± 16 | 184 ± 5 |
| **R223** | > 1000 | > 1000 | > 1000 |
| **T226** | > 1000 | > 1000 | 782 ± 130 |
| **Q249** | > 1000 | > 1000 | > 1000 |
| **R252** | 80 ± 17 | 96 ± 8 | 172 ± 17 |
| **K256** | 444 ± 50 | 274 ± 14 | 263 ± 27 |
| **K257** | 99 ± 7 | 133 ± 7 | 384 ± 42 |
| **A258** | 174 ± 18 | 208 ± 7 | 309 ± 27 |

**Model for the role of En2-GAG interaction in retinal axon guidance**

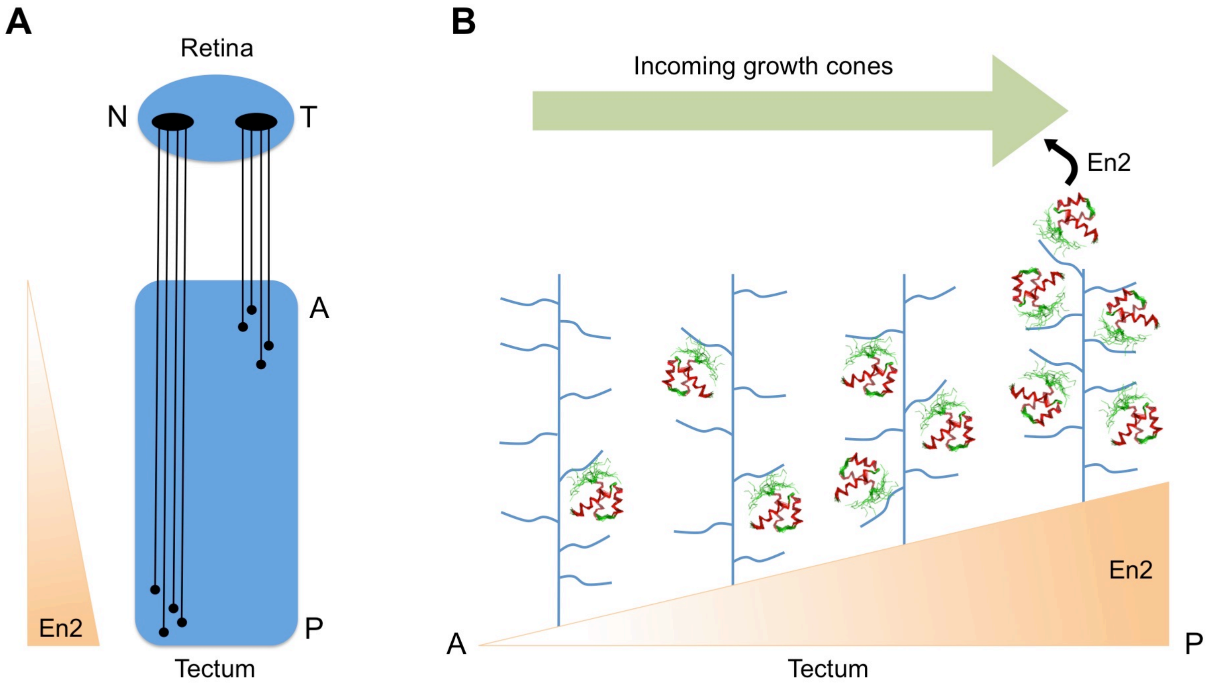

**Fig. S13**. En2 signaling in the retinotectal system. (*A*) Schematic representation of the retinotectal system showing the projection of nasal (N) and temporal (T) axons onto the anteroposterior axis of the optic tectum (A, Anterior; P, posterior). Within the tectum, En2 and Ephrin A5 morphogens show an anterior-to posterior rising gradient that is required for proper positioning of nasal and temporal axons. (*B*) Role of En2 graded expression in retinal axon guidance. En2 secreted from the tectum is maintained in the extracellular matrix at a concentration reflecting the secretion gradient, likely through electrostatic interactions with sulfated GAGs at the cell-surface that ensure a local confinement of the protein. Once internalized by growth cones in the posterior tectum, En2 stimulates the synthesis of several translational regulators, which results in Ephrin-mediated collapse of temporal growth cones.
